# Ayurveda based phenotype annotation integrated with Expectation Maximization algorithm reveal six biological clusters of rare diseases: potential in resolving ciliopathies

**DOI:** 10.1101/2024.09.13.612844

**Authors:** Aditi Joshi, Ashish Sharma, Deepika Jangir, Tanay Anand, Hemendra Verma, Manvi Yadav, Nupur Rangani, Pallavi Joshi, Ravi Pratap Singh, Sandeep Kumar, Shipra Girdhar, Rakesh Sharma, Abhimanyu Kumar, Lipika Dey, Mitali Mukerji

## Abstract

The heterogeneity of underlying causes and the variability of symptoms across multiple organs in rare diseases present significant challenges when approached from an organ-centric perspective. Ayurveda, however, takes a multisystem perspective, assessing diseases through kinetic (*Vata:V*), metabolic (*Pitta:P*), and structural (*Kapha:K*) dimensions, each with distinct phenotypic and molecular correlates. This study explores rare diseases from a systems perspective by integrating Ayurveda and unifying terminologies from both disciplines using Human Phenotype Ontology (HPO). Experts categorized 10,610 HPO terms into Ayurvedic phenotypic groups (*V/P/K*) and applied the Expectation Maximization (EM) algorithm to cluster 12,678 diseases. This yielded six distinct clusters, termed “AyurPhenoClusters,” with 2,814 diseases uniquely classified and enriched for *V/P/K* phenotypes. Functional annotation of these, highlighted key biological processes: (i) embryogenesis and skeletal morphogenesis, (ii) endocrine and ciliary functions, (iii) DNA damage response and cell cycle regulation, (iv) inflammation and immune response, (v) immune function, hemopoiesis, and telomere aging, and (vi) small molecule metabolism and transport. Notably, the *K* predominant cluster had the highest ciliary gene enrichment (43%), followed by the *V* predominant cluster (16%), suggesting potential ciliopathies in *V* cluster. This systems-based approach enhances the understanding of rare diseases by bridging Ayurveda with modern medicine, for improved diagnostics and therapeutics.

## 1 Introduction

Rare diseases characterized by complex symptoms, lack of definitive diagnostics, and limited treatment options present substantial challenges in healthcare[1]. Also, the treatments are often organ-specific, symptom centric and do not address systemic health issues [2, 3]. Large scale genome sequencing and genotype-phenotype correlations in rare diseases now reveal extensive overlap of clinical and molecular features that are gradually erasing the boundaries between seemingly unrelated diseases [4]. This highlights the need for better disease classification methods that delineate genotype to phenotypic variations more accurately for improved diagnostic yields and therapeutic outcomes [5].

Ayurveda in contrast to modern medicine defines diseases from a system’s perspective through examination of perturbation of three physiological entities, called “*tridoshas*” that govern the kinetic (*V*), metabolic (*P*) and structural attributes (*K*) of the system [6, 7]. These imbalances affect multiple body systems and are discernible by the phenotypic attributes governed by each *dosha*. Healthy individuals are stratified into 7 distinct constitution types “*Prakriti* ” on the basis of relative proportions of *V, P, K*. These constitution types differ with respect to mulitsystem attributes that include anatomical features, physiological responses as well as mental and behavioral attributes. Individuals with predominance of any one *dosha* are more prone to certain diseases and comprise 10% of the population. The diseases are also described on the basis of perturbations of *doshas* from their homeostatic thresholds. Ayurvedic treatment aims to bring *doshas* to their homeostatic thresholds through a holistic approach of drugs, diet and lifestyle modulation keeping in context the individuality of the person, the clinical history of the disease and the diseased individual [7, 8]. Ayurveda identifies more than 140 phenotypic attributes originating from a single *dosha* imbalance, each representing a simplified classification of diverse symptoms. These are described as ”*Nanatmaj Vikaras*” and herein referred as n*V*, n*P* and n*K* depending on the dosha involved. For example, pain, inflammation and obesogenic phenotype, regardless of its location in the body or general conditions, are each categorized under broad phenotypic groups of n*V*, n*P* and n*K* respectively. Research spanning over two decades has provided molecular evidence of *V*,*P*,*K* at various levels of cellular hierarchy in healthy and diseased individuals [6]. These differences map to axes of immune response, cell proliferation, inflammation, hemostasis, wound healing etc. Genetic susceptibility to complex diseases and responsiveness to stress, UV, hypoxia as well as drugs have also been linked to *doshas* [9]. We hypothesized that the Ayurvedic concepts of doshic imbalances can provide a novel framework for understanding heterogeneity seen in rare diseases, when integrated with the standardized phenotypic vocabulary of the Human Phenotype Ontology (HPO) [10, 11]. This was built on multiple lines of evidence that demonstrate distinct functional enrichments of the doshas [6, 12] and we reasoned this approach could have the potential to simplify the complexity in disease categorization, leading to more targeted and stratified interventions.

Our initial exploratory analysis with a few *dosha*-specific phenotypic groups in the Human Phenotype Ontology (HPO) provided an insight for a more extensive exercise. For instance, syndromes associated with bleeding that is a n*P* feature were predominantly enriched in *P* -related phenotypes, whereas ataxic syndromes were linked to features that more commonly associate with *V* (see Supplementary Pilot Study). With the help of domain experts, we assigned each of the 10610 HPO terms with a *dosha* label, as well as identified the HPOs that best described 140 ”*Nanatmaj Vikaras*” (n*V*, n*P* and n*K*). 9269 syndromic diseases, each of which had at least one HPO identified as (n*V*, n*P* and n*K*), were then EM clustered using the *dosha* labels assigned to the HPO terms. This led to six distinct “AyurPhenoClusters”, each of which contained diseases enriched in distinct phenotypes and biological processes. We demonstrate the utility of this clustering in identification of syndromes/diseases that have a ciliary dysfunction either due to structural or functional disruption. This first of its kind of framework promises to (a) reconcile the gap between organ-centric and system-level medical perspectives (b) provide an assistive tool to Ayurveda practitioners for clinical assessment of modern diseases and (c) provide a framework for unifying rare diseases that have shared molecular endophenotypes for enhanced management and drug repurposing.

## 2 Results

### 2.1 Mapping of HPO terms to *V* /*P* /*K* phenotypes

The pilot study of integrating Ayurveda based clinical descriptions with modern medical terminologies using the interface of HPO revealed the potential of this approach in bridging the ontological links (see Supplementary Pilot Study). Subsequently all 10,610 HPO terms associated with 12,678 diseases were annotated with *V*,*P*,*K*, and n*V*, n*P*, n*K* labels. 82% of them mapped to *V*, 11% to *P* and 7% to *K* (see Supplementary Table S1).Supplementary Table S2 shows the distribution of these labels for all the diseases. All *Nanatmaj* features with exception of a few could be mapped to one or more HPO terms (see Supplementary Table S3). Since n*V*, n*P*, n*K* features denote extreme and distinguishing perturbations of *V*,*P*,*K*, 7135 diseases which had at least one of them are selected for further analysis.

### 2.2 EM clustering reveals six clusters with distinct phenotypic and functional attributes

Using the label distributions of the 7135 diseases from Supplementary Table S2, Expectation Maximization (EM) clustering algorithm yielded six distinct AyurPhenoClusters named as C0-C5. The distribution of the diseases to the different clusters based on their memberships, are shown in Fig.1a. 2814 diseases, shown at the core, denote the diseases with membership values 1 to a cluster, and 0 to others. 3857 have a high relative membership of 0.8 to a single cluster, shown in the lighter shaded region. Diseases at the intersection have equal memberships to two clusters. Only 14 diseases have membership to more than two clusters. Strikingly, C2 stood out as an isolated cluster with no overlaps.

**Fig. 1.**
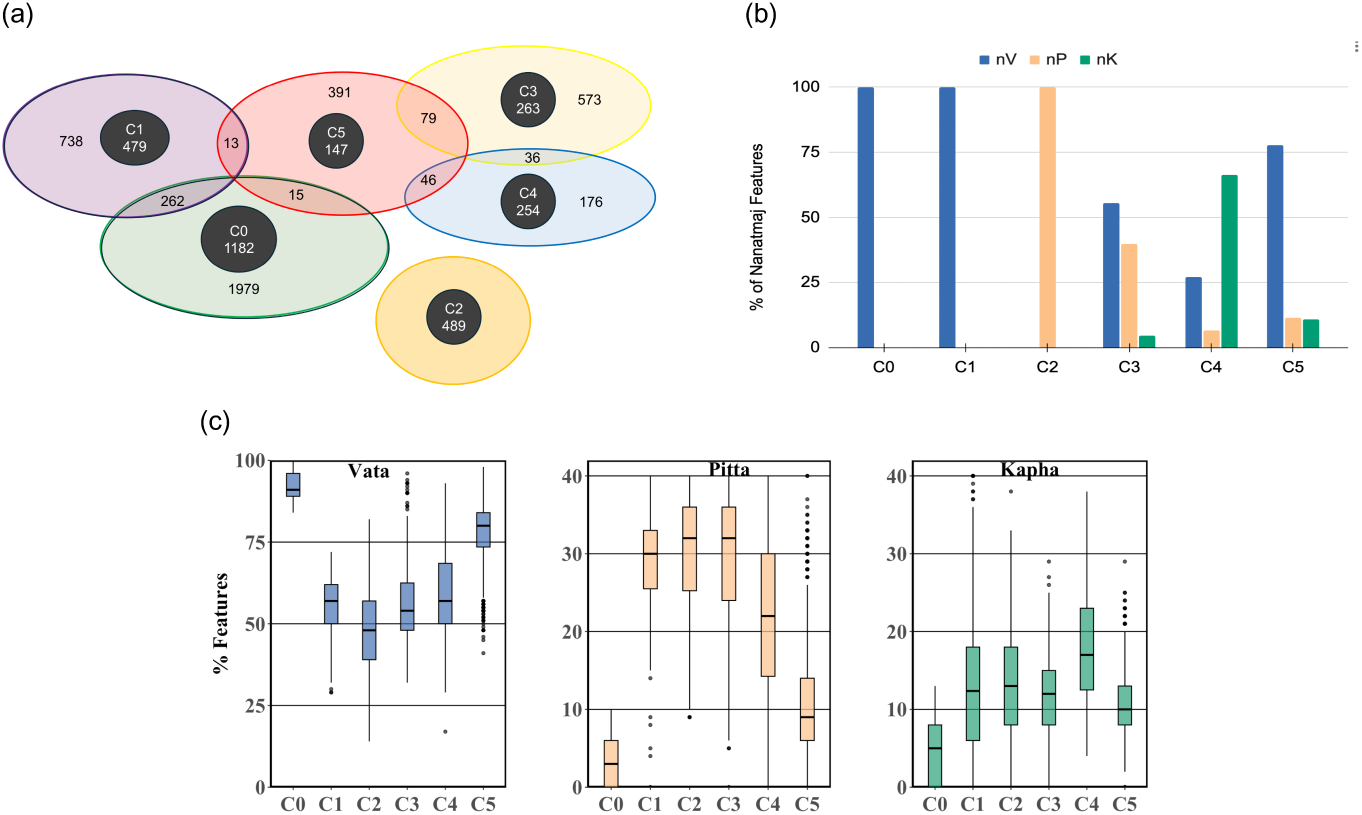
Distribution of 7135 Diseases & associated HPO features assigned to V,P,K across clusters from 2841 core diseases (a) Distribution of diseases across AyurPhenoClusters. The darker core shows number of diseases with full membership (1.0) to a cluster. The lighter region denotes diseases with majority (0.8) membership to a cluster. Diseases at intersection have equal membership to both clusters. (b) The percentage of phenotypes classified as *Nanatmaj Vikaras* for each cluster corroborates with relative enrichment of *V, P* and K features. In Cluster C0 and C1, nV is nearly exclusive whereas C2 has nearly 100% n*P*. Clusters C3, C4, and C5 have relatively lower proportions of n*V* and n*P* and nearly equal proportions of n*K*. (c) The three panels of boxplots depict the percentage of HPO terms that map to *V, P* and *K* features across each cluster. The *V* features are highest in cluster C0 and C5; *P* is highest in C3 and C2; and *K* in C4 making them predominantly *V, P* and *K* enriched clusters. Whereas C1 and C3 seem mixed *PV* types.

A deeper analysis was carried out for the 2814 diseases that mapped uniquely to a cluster (see Supplementary Table S4). A clear distinction amongst the clusters emerges from the percentage of n*V* /n*P* /n*K* and *V* /*P* /*K* features of the diseases mapped to each cluster (Fig.1b-c). Distinguishing set of features that are frequently observed in diseases of one cluster, but not as frequent in others, were identified (Table 1). Each of the six clusters is described below.

**Table 1.**
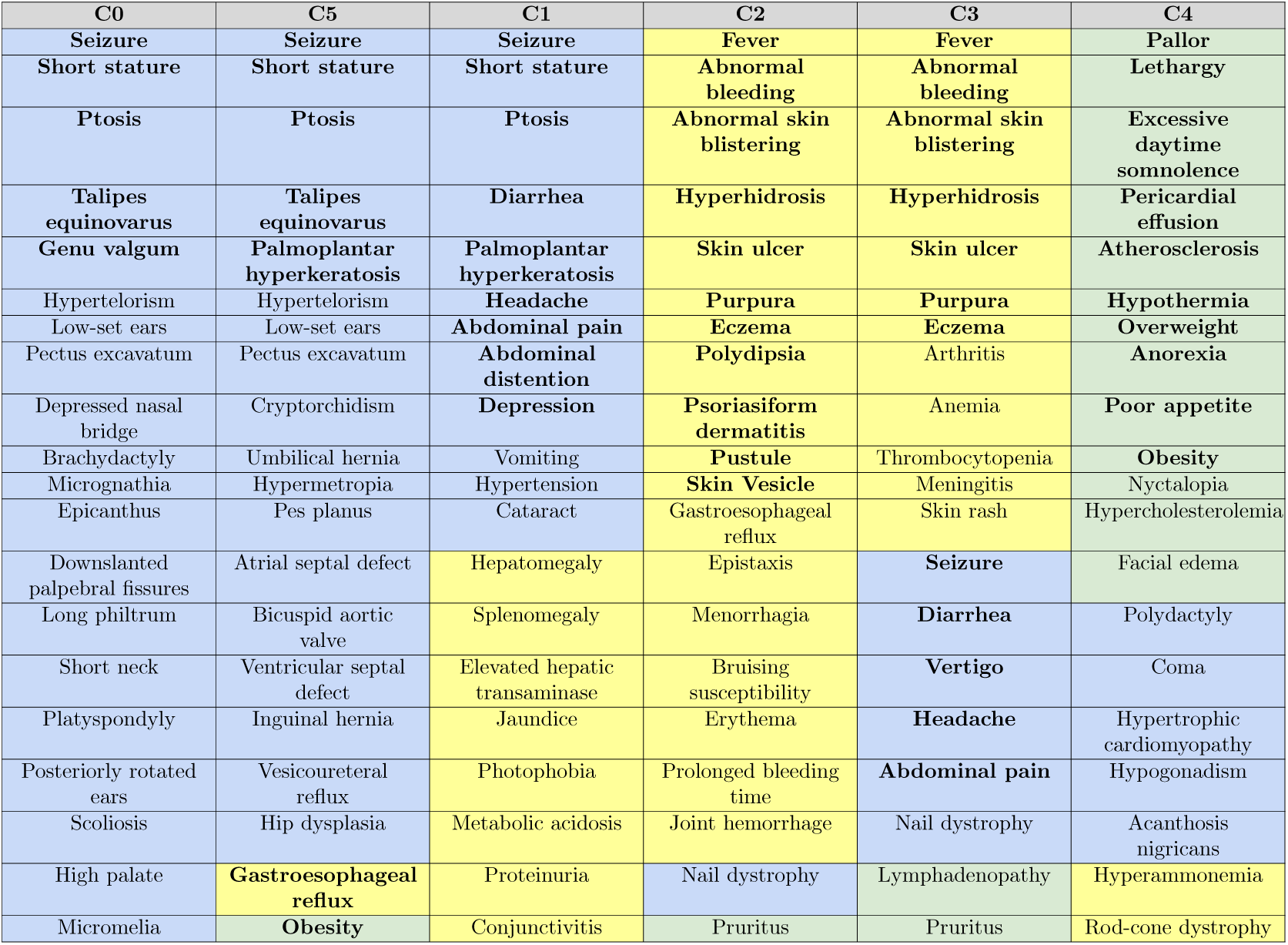
Top 20 phenotypes from the six clusters. The *V*, *P*, *K* features are colored as blue, yellow, and green, respectively. *Nanatmaj* features are underlined.

| C0 | C5 | C1 | C2 | C3 | C4 |
| --- | --- | --- | --- | --- | --- |
| Seizure | Seizure | Seizure | Fever | Fever | Pallor |
| Short stature | Short stature | Short stature | Abnormal bleeding | Abnormal bleeding | Lethargy |
| Ptosis | Ptosis | Ptosis | Abnormal skin blistering | Abnormal skin blistering | Excessive daytime somnolence |
| Talipes equinovarus | Talipes equinovarus | Diarrhea | Hyperhidrosis | Hyperhidrosis | Pericardial effusion |
| Genu valgum | Palmoplantar hyperkeratosis | Palmoplantar hyperkeratosis | Skin ulcer | Skin ulcer | Atherosclerosis |
| Hypertelorism | Hypertelorism | Headache | Purpura | Purpura | Hypothermia |
| Low-set ears | Low-set ears | Abdominal pain | Eczema | Eczema | Overweight |
| Pectus excavatum | Pectus excavatum | Abdominal distention | Polydipsia | Arthritis | Anorexia |
| Depressed nasal bridge | Cryptorchidism | Depression | Psoriasisiform dermatitis | Anemia | Poor appetite |
| Brachydactyly | Umbilical hernia | Vomiting | Pustule | Thrombocytopenia | Obesity |
| Micrognathia | Hypermetropia | Hypertension | Skin Vesicle | Meningitis | Nyctopia |
| Epicanthus | Pes planus | Cataract | Gastroesophageal reflux | Skin rash | Hypercholesterolemia |
| Downslanted palpebral fissures | Atrial septal defect | Hepatomegaly | Epistaxis | Seizure | Facial edema |
| Long philtrum | Bicuspid aortic valve | Splenomegaly | Menorrhagia | Diarrhea | Polydactyly |
| Short neck | Ventricular septal defect | Elevated hepatic transaminase | Bruising susceptibility | Vertigo | Coma |
| Platyspondyly | Inguinal hernia | Jaundice | Erythema | Headache | Hypertrophic cardiomyopathy |
| Posteriorly rotated ears | Vesicoureteral reflux | Photophobia | Prolonged bleeding time | Abdominal pain | Hypogonadism |
| Scoliosis | Hip dysplasia | Metabolic acidosis | Joint hemorrhage | Nail dystrophy | Acanthosis nigricans |
| High palate | Gastroesophageal reflux | Proteinuria | Nail dystrophy | Lymphadenopathy | Hyperammonemia |
| Micromelia | Obesity | Conjunctivitis | Pruritus | Pruritus | Rod-cone dystrophy |

C0 with 1182 diseases is exclusively characterized by n*V* with 80% of the HPO terms annotated as *V*. The number of HPO terms associated with each disease range between 20 and 30 with anatomical as the most prevalent phenotypes. C1, with 479 diseases, had n*V* and *V* as the most common, although *P* /*K* are also present in minor fractions. This cluster displays a diverse range of phenotypes, not dominated by any specific anatomical features. C2, with 489 diseases, is distinctly defined by n*P* features and also the highest proportions of *P* features. The features are predominantly associated with inflammatory conditions affecting the skin and bones, alongside symptoms such as excessive sweating (hyperhidrosis) and increased thirst (polydipsia).C3, with 263 diseases, is also n*P* and *P* enriched. In addition to features shared with C2, it harbors a subset of *V* -related phenotypes, including vertigo, headaches, diarrhea, and abdominal pain. C4, with 254 diseases, stands out with the highest percentage of n*K* among all clusters and a significant presence of *K* features in these diseases. Lastly, C5, with 147 diseases, is very heterogeneous in terms of *V* /*P* /*K* with higher average number of HPO terms compared to other clusters. The diseases in this cluster also show a high prevalence of n*V*, alongside an equal representation of n*P* and n*K* features. 55% of the features are classified as *V*, the highest number of any single *dosha* across all clusters.

### 2.3 Functional enrichment of AyurPhenoClusters

Functional annotation of the genes associated with the 2814 diseases from the six clusters revealed some striking differences in enrichment with respect to the molecular functions, biological processes and pathways (Fig.2). The genes within each cluster were also highly networked as evident from Metascape enrichments (see Supplementary Fig.S1a-e).

**Fig. 2.**
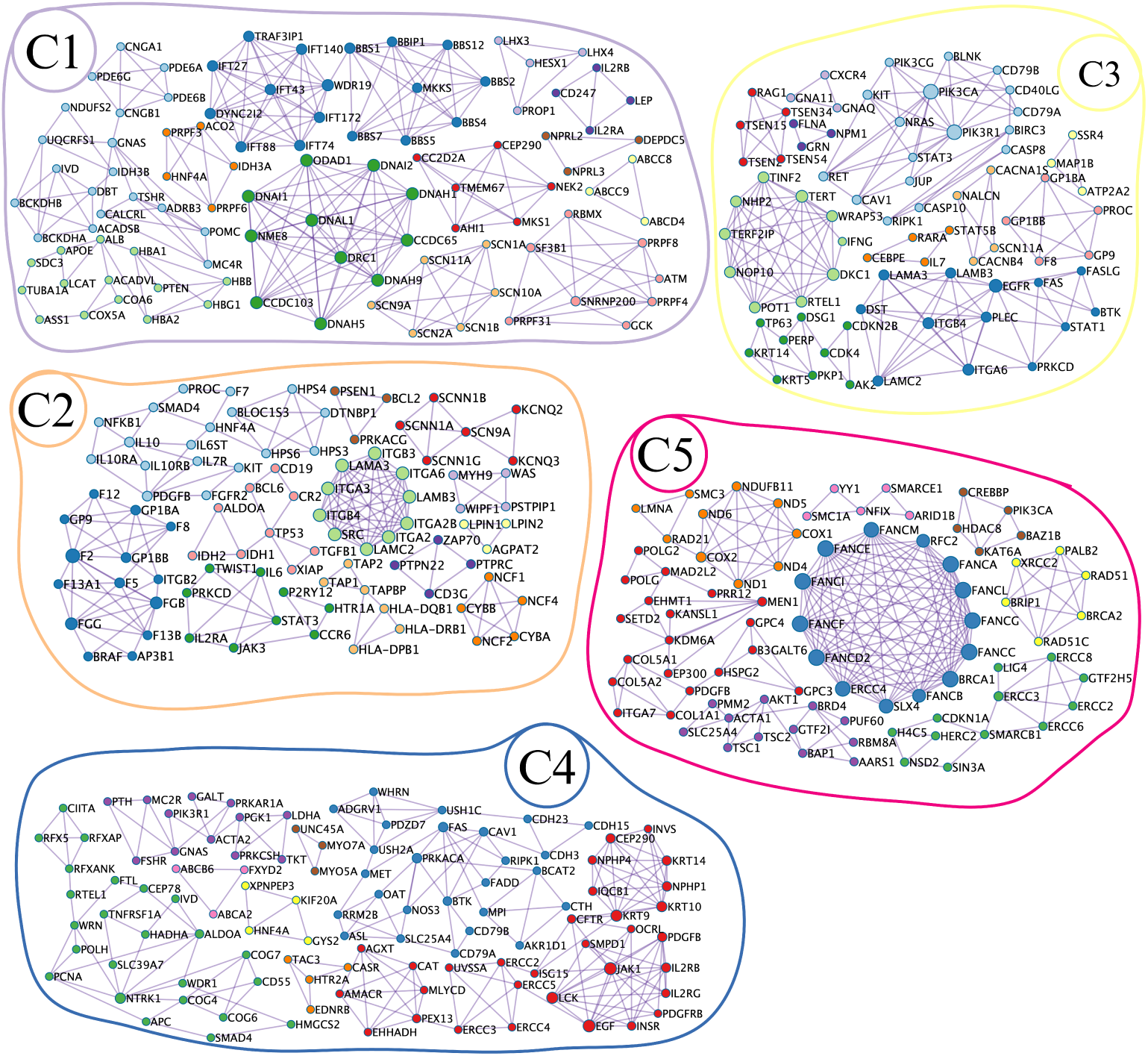
Gene enrichment network representing the EM clusters based on the distribution of HPO phenotypes associated with VPK. Nodes are colored by enriched biological and functional processes (see Supplementary Fig. S1a–e). For example, C2 (*n* = 489), predominantly *P*, is enriched with gene modules corresponding to hemopoiesis (dark brown), platelet formation (light blue), blood coagulation (dark blue), and immune response/inflammation. Cluster C4 (*n* = 254), dominated by *K*, exhibits modules related to ciliopathies (red, dark blue), transport and signaling (orange, yellow), and hormone, nutrient, and metabolic regulation. Cluster C5 (*n* = 147), with *V* features, is enriched in DNA repair modules (blue, green, yellow, brown, pink). Cluster C1 (*n* = 489) shows mixed processes such as homeostasis (yellow, blue, pink), immune (green), and signaling. Cluster C3 (*n* = 263), more *P*-like, involves hemopoiesis and telomere activity. No enriched functional modules were observed in C0 (*n* = 1182).

The *V* predominant C0 cluster was significantly enriched in processes that govern anatomy from morphogenesis, differentiation to growth and development from embryo to adult. These included development of head, limb, skeletal system and sensory organ, development, cilium assembly, bone and muscle structure. It was also enriched in processes related to regulation of cell cycle processes, membrane potential, smoothened signaling pathway, ossification and response to growth factor.

The cluster C1 was significantly enriched in processes related to cellular homeostasis, metabolism, primary immunodeficiency and renal system development. It also had processes related to response to small molecules, inorganic substances like nitrogen compounds, organic cyclic compounds as well as mechanical and light stimulus. The SHP2 pathway that regulates important cellular pathways that control proliferation, migration, differentiation, growth, and survival was also enriched in this cluster. This cluster resonates with Ayurveda description of phenotypes associated with metabolism (*P*) and cellular processes (*V*) and could be described as a *Vp* cluster.

The *P* specific C2 cluster had enrichment of processes linked to regulation of body fluid levels, immune and inflammatory processes, cell activation, cytokine signaling and production, platelet activation and TH17 cell differentiation. The associated processes are significantly perturbed in DMD, lung fibrosis, cancer especially hematopoietic lineages and govern ECM organization, multi organism level homeostasis. Heightened inflammation is a characteristic of *P* as described in Ayurveda (see Supplementary Pilot Study).

The C3 cluster had overlaps with C2 with respect to many P characteristics but also had more significant enrichment of hemopoiesis and myeloid cell differentiation, type 1 hemidesmosome assembly, telomere extension by telomerase, programmed cell death as well as response to viral and bacterial infection. Based on the phenotypic attributes, this had *Pv* features.

The *K* enriched C4 cluster was highly significant in genes related to ciliary functions as well as visual perception, determination of left right asymmetry and response and regulation of hormone, nutrient levels and metabolism of vitamin and cofactors especially Vitamin B12. These are prominent in ciliopathies, retinal dystrophy and Bardet Biedl syndrome.

Cluster C5 has an abundance of all *V*,*P*,*K* features with a predominance of *V* features. The enriched processes include DNA repair, DNA damage response, chromatin organization and regulation of cell cycle as well as response to radiation, oxygen levels. Also significant was chordate embryo development, heart development, transcriptional regulation by TP53, aminoacyl biosynthesis and protein localization and associated with diseases like Fanconi anemia, diabetic cardiomyopathy and nucleic acid metabolism and innate immune sensing.

### 2.4 AyurPhenoClusters link many rare diseases to ciliopathies

Propelled by our observation that a collection of phenotypes labeled as *K*, defining the C4 cluster, showed an enrichment in ciliopathies, we decided to investigate the potential relationship between AyurPhenoClusters and ciliopathies. We observed that 385 genes associated with different Ayurphenoclusters were present in two prominent ciliary databases, the Ciliacarta(12) and Ciliaminer, (13) and were significantly enriched in C4 (Fig.3a, see Supplementary Table S5). Also 43% of the genes in C4, 16% of the genes in C0, 11.8% of C1, 9.1% of C5, 7.5% with C3 and 5.6% in C2 were associated with ciliary structure and functions.

**Fig. 3.**
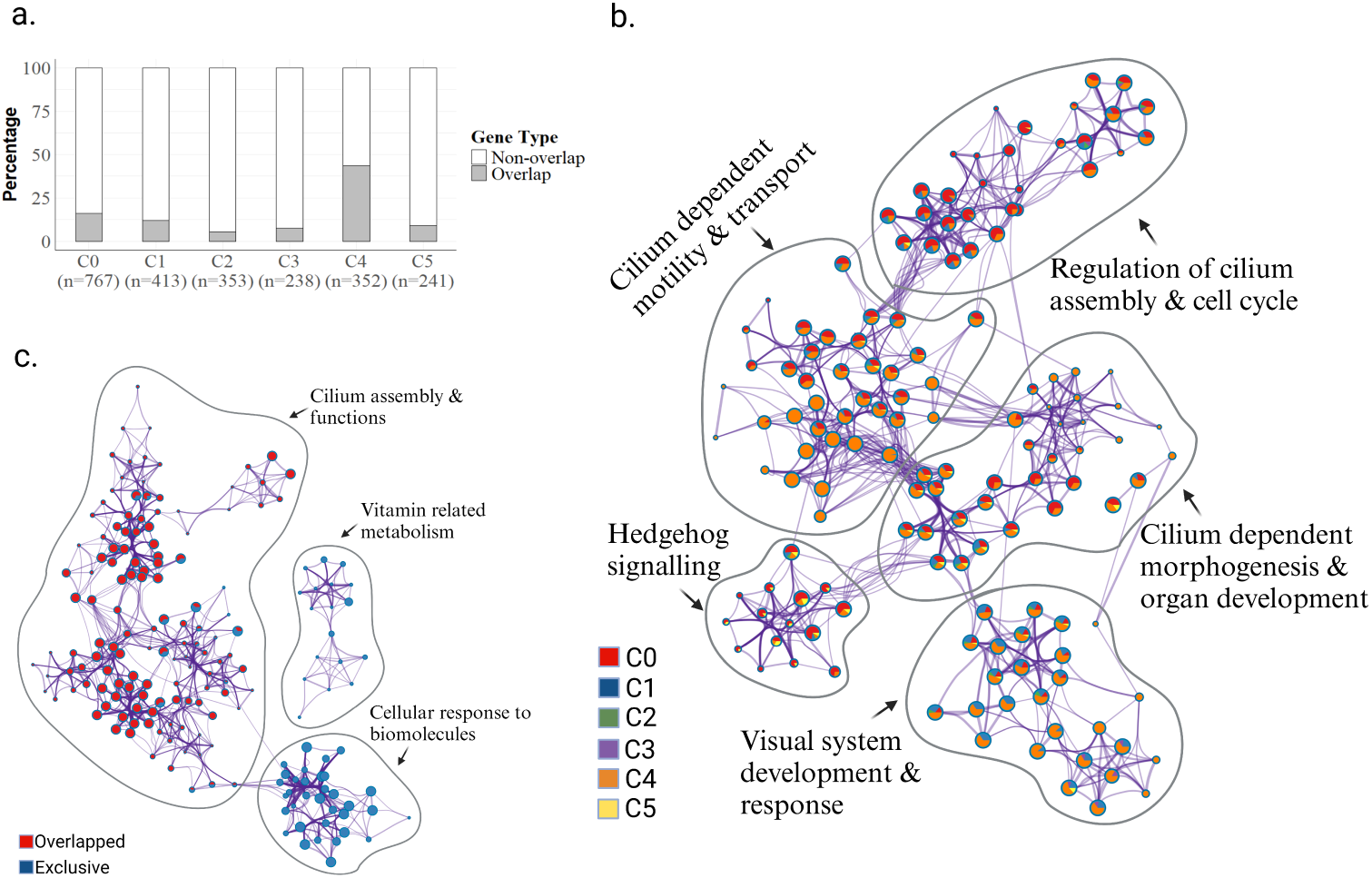
Network analysis of ciliary genes across clusters. (a) Bar plot showing the overlap between cluster genes and known ciliary genes; C4 shows the highest overlap. (b) Network analysis of ciliary genes in clusters C0–C5 reveals functional modules, including cilium-dependent morphogenesis (e.g., visual system development), cilium motility, transport, signaling, and cell cycle. Node pie charts indicate the fraction of genes from C4 (see Supplementary Fig. S4 for detailed annotation). (c) Network analysis of all genes in cluster C4 (both overlapping and exclusive) identifies three main modules, most related to cilia. The exclusive C4 set maps to cellular responses with connections to the cilia module.

Analysis using Metascape revealed extensive networks amongst the ciliary genes across all clusters (Fig.3b, see Supplementary Fig.S2). The processes within these ciliary genes are mapped to five major distinct functional hubs. These include network of genes related to (i) response to light, eye development, (ii) hedgehog signaling, embryonic development & cilium dependent cell mobility, (iii) centrosome & cell cycle regulation, (iv) regulation of cell projection & organelle assembly and (v) pathways associated with intraflagellar transport, cilium assembly, microtubule-based movement & dynein complex. There were four additional nodes that were mapped to pathways associated with protein localization. In the network, genes from C4 are prominently represented across all nodes, except those related to hedgehog signaling, embryonic development, and centrosome and cell cycle regulation in which genes from C0 have a greater contribution.

A conjoint analysis of two sets of C4 cluster genes, one mapping to reported ciliopathies and the second one comprising remaining genes, revealed three major hubs (Fig.3c, see Supplementary Fig.S3a). Two of these hubs were connected, while the third, distinct hub was associated with vitamin metabolism. Among the connected nodes, one arm was specifically related to ciliary structure and function, majorly overlapping with the reported genes. Interestingly, the other arm, which mapped to the non-overlapping set, was associated with the regulation and response to large molecules (sugars, proteins, and hormones) within the same network. A similar conjoint analysis of the network from C0 with overlapped and non-overlapped genes from the ciliary dataset revealed three distinct networks. A predominant fraction is associated with embryonic development, cell cycle and muscle movement (see Supplementary Fig.S3b). The overlapping genes from the ciliary databases are only contributing to a small part of the cluster networks associated with cilium assembly, intraflagellar transport and hedgehog signaling. Gene ORGANizer analysis reveals strikingly distinct organ enrichment of ciliary genes associated with C0 and C4 clusters (Fig.4a-b). C4 genes were highly enriched in the organs associated with endocrine, reproductive, urinary and respiratory systems whereas the expression of C0 genes were pervasive affecting skeletal, muscular and integumentary systems (see Supplementary Table S6).

**Fig. 4.**
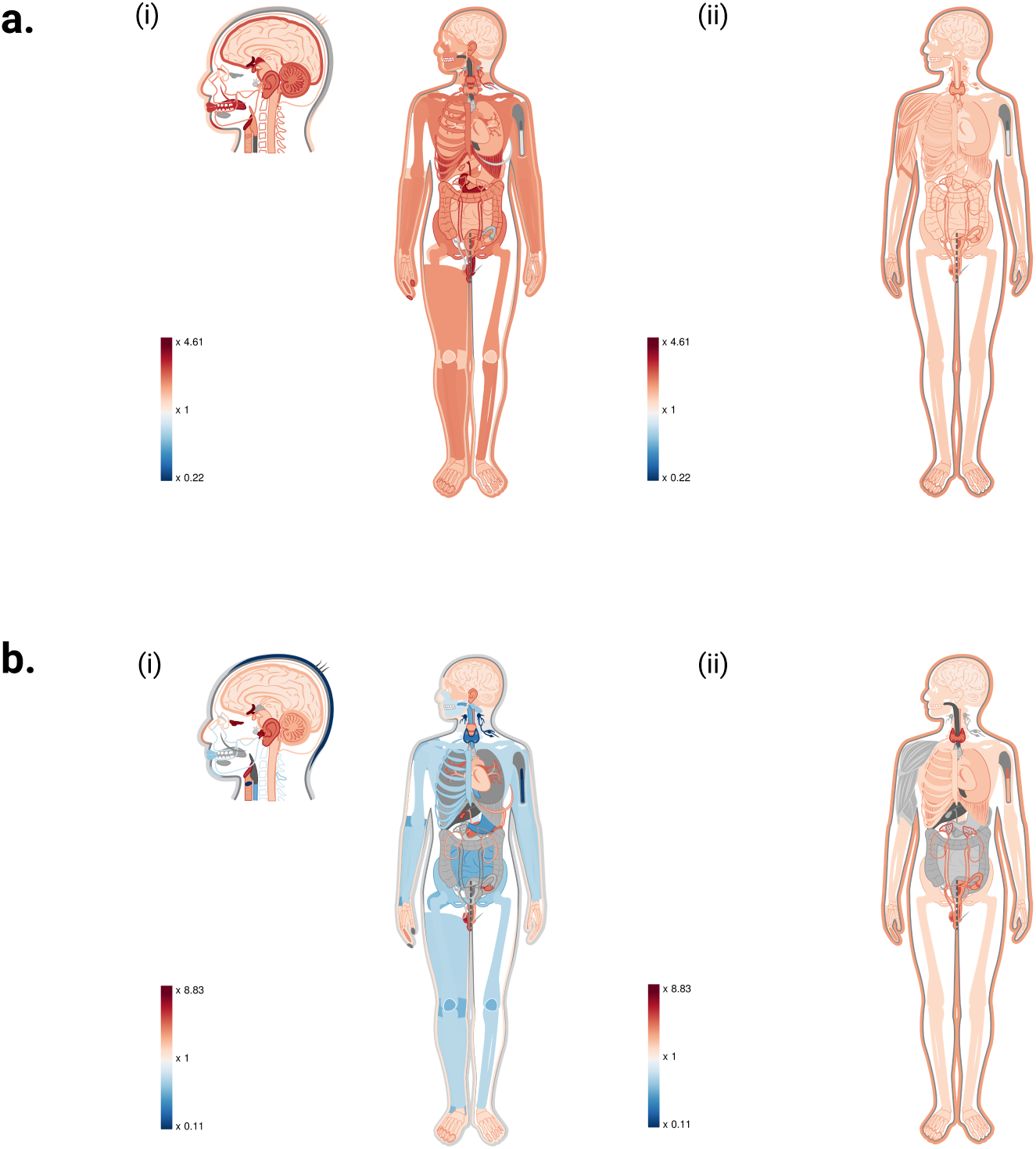
Organ-wise heatmap showing the enrichment of ciliary genes from clusters C0 and C4. Red shading indicates higher enrichment, while blue indicates depletion in the respective organs/systems. (a) Heatmap showing phenotypic expression of ciliary genes from cluster C0: (i) organ-wise and (ii) system-wise. (b) Heatmap showing phenotypic expression of ciliary genes from cluster C4: (i) organ-wise and (ii) system-wise.

### 2.5 Potential ciliopathies identified through AyurPhenoClusters

Across all the clusters, 251 diseases harbor genes reported in ciliary datasets described above. However, only 38% (95 out of 251) are reported as ciliopathies in Ciliaminer. We propose that amongst the remaining 156 diseases some could be classified as ciliopathies as (1) they share an extensive set of phenotypes with the reported ciliopathies (2) their genes are also members of the networks with ciliary structure and function. Based on Jaccard’s distance between phenotypes of reported ciliopathies and this set, we identified a few cases with high potential which include Robinow syndrome autosomal dominant 2 (OMIM:180700), VissersBodmer syndrome (OMIM:619033), Griscelli syndrome type 1 (ORPHA:79476), progressive familial intrahepatic cholestasis (ORPHA 172) etc (Table 2). We also propose that the diseases whose genes do not map to ciliary datasets but have shared memberships in AyurPhenoClusters and are well connected to the ciliary networks (Fig.3c, see Supplementary Fig.S3b) could also be due to ciliary dysfunctions. These include examples Obesity (OMIM;601665); Leptin deficiency or dysfunction (OMIM:614962); Macrocephaly/Autism syndrome (OMIM:605309); Choanal atresia and lymphedema (OMIM:6133611) etc. As is evident from Table 2, most of the potential ciliopathies are from C0.

**Table 2.** Known and potential ciliopathies across clusters. The number of known ciliopathies and potential ciliopathies were identified based on overlaps of HPO phenotypes with known ciliopathies using Jaccard similarity.

| Cluster | Known Ciliopathies (Number of Diseases) | Potential Ciliopathies (Number of Diseases Sharing Ciliary Genes) | Top 5 Diseases Based on HPO Overlaps with Known Ciliopathies (Intersect/Union of HPO sets) |
| --- | --- | --- | --- |
| C0 | 38 | 74 | Robinow syndrome autosomal dominant 2; Vissers-Bodmer syndrome; Mosaic variegated aneuploidy syndrome 2; Lissencephaly 4; Robinow syndrome, autosomal dominant 1 |
| C1 | 3 | 26 | Microcephaly and chorioretinopathy, autosomal recessive 1; Griscelli syndrome, type 1; Lipodystrophy, congenital generalized, type 3; Cholestasis, progressive familial intrahepatic, 10 |
| C2 | 9 | 16 | Crisponi/cold-induced sweating syndrome 3; Bleeding disorder, platelet-type 19; Diabetes mellitus, insulin-dependent 1; Pick disease of brain; Carney complex, type 1 |
| C3 | 0 | 21 | Deafness, neurosensory, autosomal recessive 2; Cholestasis, progressive familial intrahepatic 9; Griscelli syndrome, type 2; Episodic ataxia, type 5; Autoimmune disease, multisystem, infantile-onset 1 |
| C4 | 38 | 7 | Retinal dystrophy and obesity; Renal cysts and diabetes syndrome; Pseudohypoparathyroidism, type IB |
| C5 | 7 | 12 | Glycosylphosphatidylinositol biosynthesis defect 18; Wolf-Hirschhorn syndrome; Congenital disorder of glycosylation, type Ia; Prader-Willi syndrome |

## Discussion and Conclusion

This study presents a system-based approach to study rare genetic disorders that uses Ayurveda based characterization of HPOs and groups them into six phenotypically distinct clusters. Interestingly, many seemingly unrelated rare diseases were found to share molecular endophenotypes. We highlight the merit of this approach by demonstrating how this resolves the diagnostic dilemma of ciliopathies. The ayurvedic approach offers a holistic perspective of diseases which contrast with the focus on organ-level manifestations of modern medicine which often leaves rare disease patients with inadequate diagnostic and therapeutic options [13, 14]. This first of its kind of study not only introduces a novel approach for identifying genes associated with pathogenesis but also reveals opportunities for repurposing treatments by grouping seemingly unrelated diseases.

Earlier studies using multi-omic approaches have revealed genome-wide differences in individuals with predominance of *V*, *P* and *K* doshas. These differences have been reported across biological scales of organization viz. expression, genetic, epigenetic, metagenomic, cellular, and even physiological levels [7, 9, 15–18]. The clusters defined in this study corroborate these observations, and the differential association of these clusters to various disease classes also resonates with textual descriptions in Ayurveda (see Supplementary Pilot Study). The ability of *Nanatmaj* features to reduce dimensions in a cluster suggests that including many of these non-invasive attributes in clinical settings could further enhance stratification of a larger number of rare diseases into distinct groups. Inclusion of these features in prospective cohorts might further enhance endophenotype identification.

The use of the Expectation Maximization (EM) algorithm further enriches this study by probabilistically clustering diseases based on a flexible, data-driven approach [19]. This method effectively handles the complexity and heterogeneity inherent in large-scale genomic data, allowing for the organic discovery of disease groupings based on underlying biological patterns rather than superficial symptoms or organ systems [20, 21]. This approach reveals that conditions traditionally segregated into distinct categories may share underlying molecular mechanisms, suggesting a more interconnected biological network than previously recognized. These observations also resonate with earlier studies where unsupervised clustering using machine learning approaches as well as exome studies revealed inherent connectivity between multisystem phenotypic attributes [15, 22].

These findings have significant implications for diagnostics and therapeutic strategies in diseases exhibiting extensive clinical heterogeneity and could also become relevant in different organ specific syndromes. For instance, ciliopathies, which are often difficult to diagnose due to varied symptoms across different organs [23], were found to cluster predominantly in groups characterized by *K* or *V* features. Ciliary function plays a dual role in various diseases, acting as a cause in genetic disorders known as ciliopathies and consequential in other conditions where cilia are damaged by disease processes. Ayurveda ascribes transport and motility as functions governed by *V*, dysfunction of which leads to *K* manifestation (see Supplementary Text). In ciliopathies like primary ciliary dyskinesia and polycystic kidney disease, dysfunctional or structurally abnormal cilia directly cause the disease by impairing cellular functions and signaling pathways [24]. Conversely, in diseases such as Chronic obstructive pulmonary disease (COPD) or severe asthma, ciliary dysfunction can occur as a secondary consequence of inflammation or other cellular damage, worsening the disease’s symptoms and progression [25]. Inclusion of clinical features that define the C4 cluster with genetic testing panels might enable better resolution of ciliopathies and lead to more targeted interventions addressing specific molecular pathways active in these clusters.

These observations corroborate with the organ specific enrichment of genes from C0 and C4 clusters. C0, enriched in developmental processes could affect nearly all parts of the body and defects in these genes could result in skeletal ciliopathies. C4 ciliary genes enriched in endocrine organs could govern physiological functions that respond to external cues (hormonal and macromolecules) and when disrupted could present with diseases/phenotypes associated with the cluster. Our results highlight that a system based approach through integration of Ayurveda could not only help diagnose ciliopathies better but also resolve them into distinct endophenotypes based on doshas that could benefit from differential and more targeted treatments.

Beyond this, our approach identifies several diseases with potential ciliary involvement, with cluster C0 emerging as the largest hub. This cluster includes disorders within the centrosome–cilia axis, such as Seckel syndrome (OMIM:613823) [26, 27], as well as tubulinopathies increasingly linked to ciliopathies [28]. Interestingly, diseases not classically associated with cilia, including Alzheimer disease 3 (OMIM:607822) [29] and Epileptic encephalopathy, early infantile, 56 (OMIM:617665), also co-cluster, consistent with reports of abnormal primary cilia or cerebrospinal fluid flow defects due to dysfunction of motile cilia respectively [30]. Beyond C0, cluster C4 contains established ciliopathy phenotypes, such as retinal dystrophy and obesity (OMIM:616188) is a clinical feature in Bardet–Biedl syndrome (BBS) [31] and disorders like HNF1B-related renal cysts and diabetes (OMIM:137920), which show indirect ciliary associations [32]. These findings highlight the need for systematic validation of additional candidate diseases and suggest avenues for drug repurposing in rare disorders.

Furthermore, the cluster analysis indicated that aberrant response functions of cilia could contribute to ciliary dysfunctions in diverse diseases with endocrine, retinal, renal and skeletal diseases as the most prominent manifestations. Inflammation and DNA damage response, like ciliopathy, are core processes in various diseases, potentially being the cause and symptomatic outcome. Our analysis identified two clusters, C2 and C3, that were enriched in immune and inflammatory processes, and C5, which was distinctly enriched in DNA damage (see Supplementary Fig.S4) with distinct phenotypes (Table 1).

Our study highlights the need to redefine diseases by considering systemic phenotypic effects and molecular mechanisms, a paradigm shift from traditional organ-specific definition. This study primarily focused on a subset of rare diseases and was based on the phenotypic observations from modern medicine. Future research will focus on validating the molecular attributes of the cluster, investigating potential ciliopathies, and incorporating Ayurvedic descriptions into clinical settings. In summary, integrating Ayurvedic dosha classifications with modern genomic data through advanced computational techniques offers a compelling new perspective on rare disease classification and treatment.

## Methods

### 2.1 Annotation of HPO terms with respect to Doshas

Ayurveda classifies diseases into two broad groups based on involvement of single or multiple *doshas* as *Nanatmaj* and *Samanayaj* respectively. The phenotypes are more generic in the latter group, while specific in former(see Supplementary Pilot Study). We explored these proportions in the HPO dataset (May version, dated 2022-10-05) that had 12678 diseases and 10610 non redundant features and associated genes. The following exercise was undertaken.

### 2.2 Mapping of HPO terms to dosha attributes

A team of domain experts manually curated each of the HPO terms with respect to their correlation with either of the three *doshas* based on the foundational principles of Ayurveda as described in the ancient texts. This encompasses the entire spectrum from onset to progression to prognosis of disease. The correlation included a thorough assessment of disease mechanisms, causative factors including diet, lifestyle, and environmental influences, early warning signs, clinical symptoms at the onset, and treatment responses. Each *dosha* also bears specific attributes and deviation of phenotypic expressions from a baseline health during states of imbalance. For instance, attributes of dryness are a hallmark of *V* imbalance, manifesting across multiple organ systems as symptoms like dry skin (HP:0000958), dry mouth (HP:0000217), vaginal dryness (HP:0031088), dry cough (HP:0031246), and dry eyes (HP:0001097). On the other hand, an excess of *P* may present with bleeding (HP:0000421, HP:0040242) and burning sensations (HP:0032143, HP:6000420), while *K* imbalance might be reflected in symptoms of anorexia (HP:0002039), excessive salivation (HP:0003781), and lethargy (HP:0001254, HP:0011973). The detailed description of this mapping is mentioned in Supplementary Pilot Study.

### 2.3 Mapping of Nanatmaj Vikaras to HPO terms: clinical aspects

The experts referred to several textual references from Ayurveda literature [33] to match descriptions of diseases, or ”*Vikaras*,” from Ayurveda with HPO terms that represent corresponding phenotypic features of diseases. The features were identified based on the prominence of the organ or the system in the textual references. There are key differences between Ayurveda and modern medicine for defining diseases. This was taken into consideration during annotation of all *Nanatmaj Vikaras* as described in Supplementary Pilot Study. Supplementary Table S3 provides mapping of all *Nanatmaj Vikaras* with the HPO terms. There were a few descriptions that did not have relevant matches.

### 2.4 Re-annotation of rare syndromes from Ayurveda perspective

Each Rare disease in HPO has multiple feature attributes. A new representation of each of the diseases was created in the *dosha* space using the HPO terms mapping to the *dosha* attributes as described in the Supplementary Pilot Study). The numbers of n*V*, n*P* or n*K* represented as percentages were also included for each of the diseases.

### 2.5 Identification of clusters of diseases based on *doshic* proportions

We used Expectation Maximization (EM) algorithm to explore whether there are distinct groups or “clusters” of diseases with similar patterns of *dosha* distributions. The EM algorithm assumes that the data is generated from a mixture of several Gaussian distributions, with each representing a cluster in the data. The goal of the algorithm is to estimate the parameters of the underlying Gaussian distributions, including their means, covariances, and mixture weights or the prior probabilities. The algorithm outputs an optimal number of clusters along with a membership value between 0 and 1 for each disease to each cluster. The detailed algorithm is explained in Supplementary Methods.

### 2.6 Identification of distinguishing features for the clusters

To identify the distinguishing features of each cluster, we introduced a novel measure that could capture the significance *σ* of a feature f for a cluster c using a formula that assigns a high value to a feature that is unique for a cluster and frequent within the cluster. For each cluster c, let *δ*(f, c) denote the percentage of diseases in the cluster that contain the HPO feature f. The significance *σ*(f, c) is calculated as below:

If x(f, c) = 0, then *σ*(f, c) = 0,

Otherwise, *σ*(f, c) = x(f, c) × i^≁^=c 1 / 1+ln(x(f,i))

It assigns a low value to features that are common across clusters. To identify the top n features of a cluster, the features are sorted in decreasing order of significance.

### 2.7 GO and network analysis of OMIM genes associated with clusters

In order to comprehensively understand the clusters at the molecular level we used Metascape. Along with functional enrichment it also performs interactome analysis and annotates the gene sets to functional modules based on information from over 40 independent knowledgebases [34]. We also used Metascape to perform comparative analysis of functional modules/gene sets between and within the clusters.

### 2.8 Analysis of AyurPhenoClusters in ciliopathy

We downloaded data from CiliaMiner and CiliaCarta databases [35, 36], both of which are valuable resources for ciliopathy-related diseases and their genes. The CiliaMiner database has more than 500 diseases with their associated genes categorized into primary, secondary, motile, and atypical, on the basis of subcellular localization and functions. Among these, 274 genes are reported as potential ciliopathy candidates, as these are involved in the formation, function, and maintenance of cilia. On the other hand, Ciliacarta has a total of 956 genes, of which 302 are from the SYSCILIA gold standard, 677 from gene ontology and 285 from Bayesian predictions. We analyzed the genes from AyurphenoClusters that intersected with the above databases.

Cluster-wise network analysis was conducted to identify all pathways associated with the overlapping genes using METASCAPE. Additionally, a conjoint analysis with METASCAPE was performed on two sets of genes from each of the two clusters (C4 and C0) enriched in ciliary genes: one set consisted of reported ciliary genes, and the other set consisted of the remaining genes in the cluster. These genes were mapped to rare diseases in the AyurPhenoClusters and phenotypically compared with those reported in CiliaMiner. This led to identification of unreported potential ciliopathies in the set of rare diseases.

To broadly understand the organ centric effect of cillary genes from two contrasting phenotypic clusters C0 (*V* dominant) and C4 (*K* dominant), we used Gene ORGANizer [37]. This helped to assess the differences in expression enrichment of these genes across different organs. This web-based tool, powered by an extensively curated database of over 150,000 gene-organ associations, links more than 7,000 genes to approximately 150 anatomical parts.

## Acknowledgements

MM, AJ and DJ acknowledge financial support from MOA (Ministry of AYUSH) for Center of Excellence ”AyurTech” (S/MOA/MT-M/AA/20210105), IIT Jodhpur. AS acknowledges the University Grants Commission (UGC), Government of India, for financial support of his PhD. Authors acknowledge IIT Jodhpur and DSRRA University for infrastructure support. The authors would also like to thank Aniket Bhattacharya for critical inputs.

## Conflict of Interest

The authors declare no competing interests.

## Ethics approval and consent to participate

Not applicable

## Consent for publication

Not applicable

## Data availability

Data is provided within the manuscript or supplementary information files and can also be accessed from https://github.com/Aditi-GenomicsLab/AyurPhenoCluster/.

## Materials availability

Not applicable

## Code availability

Not applicable

## Authors Contribution

MM conceived the study, MM and LD developed the study design; AK and RS lead the Ayurveda annotation; DJ, HV, SK, SG, MY, PJ, NR, RPS carried out HPO annotation, DJ annotated the *Nanantmaj Vikara*; MM and LD lead the data analysis; LD designed the computational methodology, and MM lead the genomics study; AJ and TA curated the diverse datasets and performed the data analysis with LD ; AS and AJ worked on ciliopathy and illustration; DJ, AK, AS, LD and MM synthesized the work; AJ, AS, LD and MM wrote the manuscript.

## Notes

https://github.com/Aditi-GenomicsLab/AyurPhenoCluster/

### Competing Interest Statement

The authors have declared no competing interest.

### Summary of Updates

The abstract has been modified. Sections like Method and Discussion have been elaborated more. Supplementary has been updated with indexing and details.

