## Supplementary Text for "Ayurveda based phenotype annotation integrated with Expectation Maximization algorithm reveal six biological clusters of rare diseases: potential in resolving ciliopathies"

**This file includes:**

1. **Pilot Study**
2. **Text**
3. **Methods**
4. **Figures S1-S4**

1. **Pilot study**

**Preliminary investigation into using HPO for integrating Ayurvedic and Modern clinical terminologies**

The ancient texts of Ayurveda provide descriptions of diseases in Sanskrit and classify the clinical conditions based on perturbations of the *Vata (V), Pitta (P),* and *Kapha (K)* doshas (1, 2). The diseases that have predominance of a single *dosha* are called *Nanatmaj Vikara* and 80,40 and 20 conditions have been ascribed to *V, P* and *K* respectively (see the **boxes 1,2 and 3** mentioned below). However, in contemporary times, Ayurveda clinicians examine patients who present with a gamut of symptoms described by modern medical terminologies. The clinician has to diagnose these diseases based on the closest matching descriptions from their texts for evolving treatment strategies.

1. ***Vattaj Nanatamja Vikaras-*** The 80 clinical conditions/phenotypes that are described for *Vata* Perturbations are translated from original sanskrit to the closest english descriptions

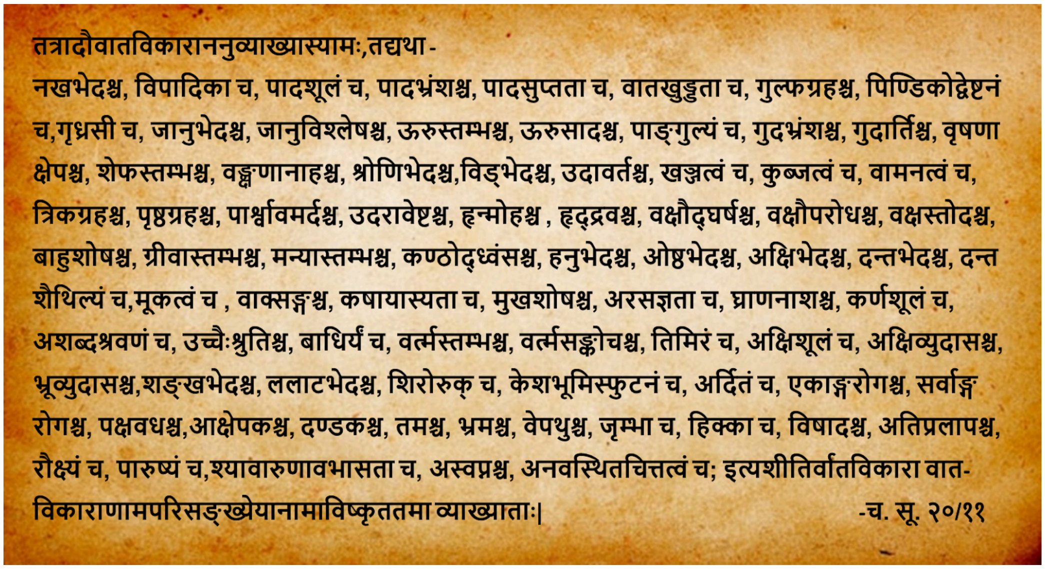
तत्रादौवातविकाराननुव्याख्यास्यामः,तद्यथा -
नखभेदश्च, विपादिका च, पादशूलं च, पादभ्रंशश्च, पादसुप्तता च, वातखुड्डता च, गुल्फग्रहश्च, पिण्डिकोद्वेष्टनं च,गृध्रसी च, जानुभेदश्च, जानुविश्लेषश्च, ऊरुस्तम्भश्च, ऊरुसादश्च, पाङ्गुल्यं च, गुदभ्रंशश्च, गुदार्तिश्च, वृषणा क्षेपश्च, शेफस्तम्भश्च, वङ्क्षणानाहश्च, श्रोणिभेदश्च,विड्भेदश्च, उदावर्तश्च, खञ्जत्वं च, कुब्जत्वं च, वामनत्वं च, त्रिकग्रहश्च, पृष्ठग्रहश्च, पार्श्वावमर्दश्च, उदरावेष्टश्च, हृन्मोहश्च , हृद्द्रवश्च, वक्षौद्घर्षश्च, वक्षौपरोधश्च, वक्षस्तोदश्च, बाहुशोषश्च, ग्रीवास्तम्भश्च, मन्यास्तम्भश्च, कण्ठोद्ध्वंसश्च, हनुभेदश्च, ओष्ठभेदश्च, अक्षिभेदश्च, दन्तभेदश्च, दन्त शैथिल्यं च,मूकत्वं च , वाक्सङ्गश्च, कषायास्यता च, मुखशोषश्च, अरसज्ञता च, घ्राणनाशश्च, कर्णशूलं च, अशब्दश्रवणं च, उच्चैःश्रुतिश्च, बाधिर्यं च, वर्त्मस्तम्भश्च, वर्त्मसङ्कोचश्च, तिमिरं च, अक्षिशूलं च, अक्षिव्युदासश्च, भ्रूव्युदासश्च,शङ्खभेदश्च, ललाटभेदश्च, शिरोरुक् च, केशभूमिस्फुटनं च, अर्दितं च, एकाङ्गरोगश्च, सर्वाङ्ग रोगश्च, पक्षवधश्च,आक्षेपकश्च, दण्डकश्च, तमश्च, भ्रमश्च, वेपथुश्च, जृम्भा च, हिक्का च, विषादश्च, अतिप्रलापश्च, रौक्ष्यं च, पारुष्यं च,श्यावारुणावभासता च, अस्वप्नश्च, अनवस्थितचित्तत्वं च; इत्यशीतिर्वातविकारा वात-विकाराणामपरिसङ्ख्येयानामाविष्कृततमा व्याख्याताः|
-च. सू. २०/११

The disorders of Vata include - cracking of soles, pain in foot, foot drop, numbness in feet, pain in ankles, stiffness in ankles, cramps in calf, sciatica, tearing pain in knees, dislocation of knees, stiffness in thighs, loss of movement in thighs, lameness, prolapse of rectum, pain in anus, twitching in scrotum, stiffness in penis, pain in groins, pain in pelvis, pain in defecation, upward movement of Vayu (gases), limping, hunch back, dwarfism, stiffness in sacral region, stiffness in back, compression in sides, twisting pain in abdomen, cardiac dysfunction, tachycardia, shivering in chest, constriction in chest, chest pain, wasting of arms, stiffness of neck, stiffness of sternomastoid, hoarseness of voice, pain in jaw, cracking of lips, pain in eyes, pain in teeth, loose teeth, dumbness, stammering, astringent taste in mouth, dryness of mouth, loss of taste sensation, loss of smelling sensation, ear-ache, dizziness in ears, hardness in hearing, deafness, stiffness in eyelids, contraction in eyelids, loss of vision, pain in eyes, squint, twisting of eyebrows, pain in the temporal region, pain in forehead, headache, cracking of scalp, facial paralysis, monoplegia, polyplegia, hemiplegia, convulsions, tetanic convulsions, feeling of darkness before eyes, giddiness, tremors, yawning, hiccup, depression, excessive delirium, roughness, coarseness, blackish and reddish luster, insomnia, instability of mind-these are the eighty most prominent ones among the innumerable disorders of Vata.

2. ***Pittaj Nanatamja Vikaras-*** The 40 clinical conditions/phenotypes that are described for *Pitta* Perturbations are translated from original sanskrit to the closest english descriptions

**पित्तविकारांश्चत्वारिंशतमत ऊर्ध्वमनुव्याख्यास्यामः-**
**ओषश्च, प्लोषश्च, दाहश्च, दवथुश्च, धूमकश्च, अम्लकश्च, विदाहश्च, अन्तर्दाहश्च, अंसदाहश्च, ऊष्माधिक्यं च, अतिस्वेदश्च(अङ्गस्वेदश्च),अङ्गगन्धश्च, अङ्गावदरणं च, शोणितक्लेदश्च, मांसक्लेदश्च, त्वग्दाहश्च, (मांसदाहश्च), त्वगवदरणं च, चर्मदलनं च, रक्तकोठश्च, रक्त विस्फोटश्च, रक्तपित्तं च, रक्तमण्डलानि च, हरितत्वं च,हारिद्रत्वं च, नीलिका च, कक्षा (क्ष्या)च, कामला च, तिक्तास्यता च, लोहितगन्धास्यता च, पूतिमुखता च, तृष्णाधिक्यं च, अतृप्तिश्च, आस्यविपाकश्च, गलपाकश्च, अक्षिपाकश्च, गुदपाकश्च, मेढ्रपाकश्च, जीवा-दानं च, तमःप्रवेशश्च, हरितहारिद्रनेत्रमूत्रवर्चस्त्वं च; इति चत्वारिंशत्पित्तविकाराः पित्त-विकाराणामपरिसङ्ख्येयानामाविष्कृततमा व्याख्याताः|**
**-च. सू. २०/१४**

The disorders of Pitta includes - heating, scorching, burning, intense burning, fuming, hyperacidity, burning in stomach and esophagus, internal burning, burning in scapular region, Pyrexia, over perspiration, foul smell in body, tearing of body parts, excessive moisture in blood(ureamic condition of blood), moistening of muscles, burning io skin, tearing of skin, thickening of skin, urticarial patches, pustules, internal hemorrhage, hemorrhagic patches, greenishness, yellowness bluishness, herpes, jaundice, bitterness in mouth, bloody smell from mouth, fetid smell from mouth, excessive thirst, loss of, contentment, stomatitis, inflammation in throat, inflammation in eyes, inflammation in anus, inflammation in penis, discharge of pure blood, fainting, green or yellow color in eyes, urine and feces- these are the prominent ones among the innumerable disorders of Pitta.

3. ***Kaphaj Nanatamja Vikaras-*** The 20 clinical conditions/phenotypes that are described for *Kapha* Perturbations are translated from original sanskrit to the closest english descriptions

श्लेष्मविकारांश्च विंशतिमत ऊर्ध्वं व्याख्यास्यामः; तद्यथा- तृप्तिश्च, तन्द्रा च,
निद्राधिक्यं च, स्तैमित्यं च, गुरुगात्रता च, आलस्यं च, मुखमाधुर्यं च, मुखस्रावश्च,
श्लेष्मोद्गिरणं च, मलस्याधिक्यं च, बलासकश्च, अपक्तिश्च, हृदयोपलेपश्च,
कण्ठोपलेपश्च, धमनीप्रति(वि)चयश्च, गलगण्डश्च, अतिस्थौल्यं च, शीताग्निता च,
उदर्दश्च, श्वेतावभासता च, श्वेतमूत्रनेत्रवर्चस्त्वं च; इति विंशतिः श्लेष्मविकाराः
श्लेष्मविकाराणामपरिसङ्ख्येयानामाविष्कृततमा व्याख्याता भवन्ति||

-च. सू. २०/१७

The disorders of *Kapha* includes- saturation, drowsiness, excessive sleep, cold sensation, heaviness in body, lassitude, sweetness in mouth, salivation, mucous expectoration, excess of dirt, excess of mucus, indigestion, plastering of heart, plastering of throat, accumulation in vessels, goiter, over plumpness, urticarial eruptions, urticarial patches, white luster, whiteness in urine, eyes and feces-these twenty are the prominent ones among the innumerable disorders of kapha.

Following the above Ayurveda descriptions in boxes 1, 2 and 3 above, some of the important considerations during mapping are highlighted below.

#### 1.1 Mapping of HPO-ID to *dosha* attributes : Clinical aspects

Each *dosha* bears specific attributes that influence the phenotypic presentation during states of imbalance. For instance, attributes of dryness are a hallmark of *Vata* imbalance, manifesting across multiple organ systems as symptoms like dry skin (HP:0000958), dry mouth (HP:0000217), vaginal dryness (HP:0031088), dry cough (HP:0031246), and dry eyes (HP:0001097). These and other related signs to dryness are thus categorized under *Vata*-related phenotypes. *Dosha* classifications were further refined by the descriptions that reflect deviation of phenotypic expressions from a person's baseline health. Elevated *Vata*, for example, is characterized by phenotypes associated with pain (HP:0012514, HP:0003418) and muscle cramps (HP:0003394, HP:0032155). On the other hand, an excess of *Pitta* may present with bleeding (HP:0000421, HP:0040242) and burning sensations (HP:0032143, HP:6000420), while *Kapha* imbalance might be reflected in symptoms of anorexia (HP:0002039), excessive salivation (HP:0003781), and lethargy (HP:0001254, HP:0011973). Also phenotypic attributes related to sweat, stool, urine etc and *dosha* involvement in symptoms from different tissues like lymph, blood, muscle, adipose, bones, marrow, reproductive tissue, skin, nervous were considered.

#### 1.2 Mapping of *Nanatmaj Vikara* to HPO IDs : clinical aspects

There are key differences between Ayurveda and modern medicine for defining diseases. This was taken into consideration during annotation of all *Nanatmaj Vikara* (denoted as "n"). For example

- Brittle nails (*Nakhbheda*): In Ayurveda, this is considered a disorder associated with the *V dosha* (n*V*), while HPO classifies it as a phenotypic abnormality of nails (HPO ID: HP:0001808) related to 28 different diseases.

- Excessive thirst (Polydipsia, *trishna-adhikya*): Ayurveda describes this as a *P*-related disorder (n*P*), whereas HPO lists it as a symptom (HPO ID: HP:0001959) found in over 62 diseases.

- Excessive sleepiness (*Nidra adhikya*): This is categorized as a *K*-related disorder (n*K*) in Ayurveda. In contrast, HPO associates it with hypersomnia (HPO ID: HP:0100786) and links it to more than 28 diseases.

To create a more objective and interoperable framework, NLP can be employed to bridge the medical terminologies of both the streams. Given that the Human Phenotype Ontology (HPO) has been integrated into various cohorts and languages (3). Utilizing HPO could connect these Ayurvedic and modern clinical descriptions, thereby expanding access to numerous existing cohort data for Ayurveda based stratification.

In order to explore the ontological links between Ayurveda and modern medical terminologies we conducted a pilot study where we explored three possibilities of linkages

1. *Nanatmaj Vikara* in HPO diseases
2. Associations of clinical features of dosha with modern diseases
3. Links to OMIM diseases from molecular links of *doshas*

**Case Study 1- *Nanatmaj Vikara* in HPO diseases**
A condition of bleeding risk (Von Willebrand disease) that is also referred to amongst one of the *Pittaj Nanatmaj Vikara* (*Raktapitta*) was first considered. In an earlier study variation in a gene VWF linked to this disease was also associated with *Pitta* phenotypes (4). A search term for "Abnormality of von Willebrand factor" in HPO yielded 19 diseases listed below (Supplementary Pilot Study Table 1) (**HPO version: May version, dated 2022-10-05**).

**<u>Supplementary Pilot Study Table 1:</u>** List of syndromes with a search for "Abnormality of von Willebrand factor" term in the HPO database

| HPO Disease Id | HPO Disease Name |
|---|---|
| ORPHA:251061 | 7q31 microdeletion syndrome |

| | |
|---|---|
| ORPHA:99147 | Acquired von Willebrand syndrome |
| ORPHA:274 | Bernard-Soulier syndrome |
| OMIM:231200 | Bernard-Soulier syndrome |
| OMIM:153670 | Bernard-Soulier syndrome, type A2, autosomal dominant |
| OMIM:614201 | Bleeding disorder, platelet-type, 11 |
| OMIM:618462 | Bleeding disorder, platelet-type, 22 |
| OMIM:619271 | Bleeding disorder, platelet-type, 24, autosomal dominant |
| ORPHA:70591 | Chronic thromboembolic pulmonary hypertension |
| OMIM:273800 | Glanzmann thrombasthenia |
| ORPHA:849 | Glanzmann thrombasthenia |
| OMIM:619267 | Glanzmann thrombasthenia 2 |
| ORPHA:79259 | Glycogen storage disease due to glucose-6-phosphatase deficiency type Ib |
| OMIM:139090 | Gray platelet syndrome |
| ORPHA:169802 | Severe hemophilia A |
| OMIM:619130 | Thrombocytopenia, autosomal dominant, 7 |
| ORPHA:903 | Von Willebrand disease |
| OMIM:193400 | von Willebrand disease, type 1 |
| OMIM:277480 | von Willebrand disease, type 3 |

Each of the above diseases are associated with many clinical features each of which are assigned an HP ID. For example one of the diseases listed above Glanzmann thrombasthenia (**OMIM ID** 273800) in HPO lists features with HP IDs that are given in Supplementary Pilot Study Table 2 below.

**Supplementary Pilot Study Table2:** List of phenotypes associated with Glanzmann thrombasthenia (**OMIM ID** 273800)

| | HPO term id | HPO term name |
|---|---|---|
| 1 | HP:0011873 | Abnormal platelet count |
| 2 | HP:0000007 | Autosomal recessive inheritance |
| 3 | HP:0000978 | Bruising susceptibility |
| 4 | HP:0001975 | Decreased platelet glycoprotein IIb-IIIa |
| 5 | HP:0031364 | Ecchymosis |
| 6 | HP:0000421 | Epistaxis |
| 7 | HP:0030138 | Excessive bleeding from superficial cuts |
| 8 | HP:0002239 | Gastrointestinal hemorrhage |

| | | |
|---|---|---|
| 9 | HP:0000225 | Gingival bleeding |
| 10 | HP:0004866 | Impaired ADP-induced platelet aggregation |
| 11 | HP:0031126 | Impaired clot retraction |
| 12 | HP:0008320 | Impaired collagen-induced platelet aggregation |
| 13 | HP:0008148 | Impaired epinephrine-induced platelet aggregation |
| 14 | HP:0003540 | Impaired platelet aggregation |
| 15 | HP:0011871 | Impaired ristocetin-induced platelet aggregation |
| 16 | HP:0002170 | Intracranial hemorrhage |
| 17 | HP:0000132 | Menorrhagia |
| 18 | HP:0003623 | Neonatal onset |
| 19 | HP:0003010 | Prolonged bleeding time |
| 20 | HP:0000979 | Purpura |
| 21 | HP:0100309 | Subdural hemorrhage |

All the HP IDs (149 features) related to the above syndromes were provided to domain experts/Ayurveda clinicians for assignment into *V, P* and K based on their clinical description and matches to Ayurveda texts. These features were then mapped back to each of the OMIM diseases. Supplementary Pilot Study Table 3 below illustrates an example of such a labeling with their corresponding mapping to Glanzmann thrombasthenia.

**Supplementary Pilot Study Table 3:** Labeling of the V/P/K phenotypes in one of the diseases, Glanzmann thrombasthenia (**OMIM ID** 273800)

| **HPO Disease Id** | **HPO Disease Name** | **HPO Term Id** | **HPO Term Name** | ***VPK* Mapping** |
|---|---|---|---|---|
| OMIM:273800 | Glanzmann thrombasthenia | HP:0000132 | Menorrhagia | *V/P* |
| | | HP:0030138 | Excessive bleeding from superficial cuts | *P* |
| | | HP:0003540 | Impaired platelet aggregation | *V/K* |
| | | HP:0003010 | Prolonged bleeding time | *V* |
| | | HP:0031364 | Ecchymosis | *P* |
| | | HP:0002170 | Intracranial hemorrhage | *P* |
| | | HP:0031126 | Impaired clot retraction | *P/K* |
| | | HP:0011873 | Abnormal platelet count | *V* |
| | | HP:0011871 | Impaired ristocetin-induced platelet aggregation | *K* |
| | | HP:0000979 | Purpura | *K/nP* |
| | | HP:0008320 | Impaired collagen-induced platelet aggregation | *V* |
| | | HP:0000007 | Autosomal recessive inheritance | *V* |
| | | HP:0100309 | Subdural hemorrhage | *P* |

| | | | | |
|---|---|---|---|---|
| | | HP:0000978 | Bruising susceptibility | *P* |
| | | HP:0008148 | Impaired epinephrine-induced platelet aggregation | *V* |
| | | HP:0002239 | Gastrointestinal hemorrhage | *P* |
| | | HP:0000421 | Epistaxis | *P* |
| | | HP:0004866 | Impaired ADP-induced platelet aggregation | *K* |
| | | HP:0001975 | Decreased platelet glycoprotein IIb-IIIa | *V* |
| | | HP:0003623 | Neonatal onset | *V* |
| | | HP:0000225 | Gingival bleeding | *P* |

The cumulative count of *V/P/K* features for each disease was aggregated in a matrix as shown below (Supplementary Pilot Study Table 4) and plotted to observe the trend of labeled phenotypes (Supplementary Pilot Study Fig.1). The results reveal significant differences in the number of *V/P/K* features. *V* features are highest (as described below in Supplementary Methods) followed by *P* and *K*. Bleeding is associated with imbalance of *P* and the proportion of *P* seems to be reflected in all the diseases associated with bleeding. This would be more evident when we compare it with the other examples below. These diseases to an Ayurveda doctor in a clinical setting would be considered as an imbalance of *Pitta* and termed as *Raktapitta* .

**Supplementary Pilot Study Table 4:** Cumulative count of phenotypes labeled as *V, P* and *K* for the diseases associated with “Abnormality of von Willebrand factor”

| **HPO Disease Id** | **HPO Disease Name** | **Total *V* Count** | **Total P Count** | **Total K Count** |
|---|---|---|---|---|
| OMIM:273800 | Glanzmann thrombasthenia | 9 | 12 | 5 |
| OMIM:619267 | Glanzmann thrombasthenia 2 | 10 | 6 | 3 |
| ORPHA:79259 | Glycogen storage disease due to glucose-6-phosphatase deficiency type Ib | 40 | 25 | 11 |
| OMIM:139090 | Gray platelet syndrome | 13 | 6 | 2 |
| ORPHA:169802 | Severe hemophilia A | 14 | 16 | 4 |
| OMIM:619130 | Thrombocytopenia, autosomal dominant, 7 | 6 | 3 | 2 |
| OMIM:193400 | Von willebrand disease, type 1 | 13 | 11 | 2 |
| OMIM:277480 | Von willebrand disease, type 3 | 10 | 10 | 1 |
| ORPHA:99147 | Acquired von Willebrand syndrome | 14 | 18 | 2 |
| OMIM:231200 | Bernard-Soulier syndrome | 11 | 9 | 2 |
| OMIM:619271 | Bleeding disorder, platelet-type, 24, autosomal dominant | 6 | 6 | 3 |
| ORPHA:70591 | Chronic thromboembolic pulmonary hypertension | 28 | 9 | 16 |

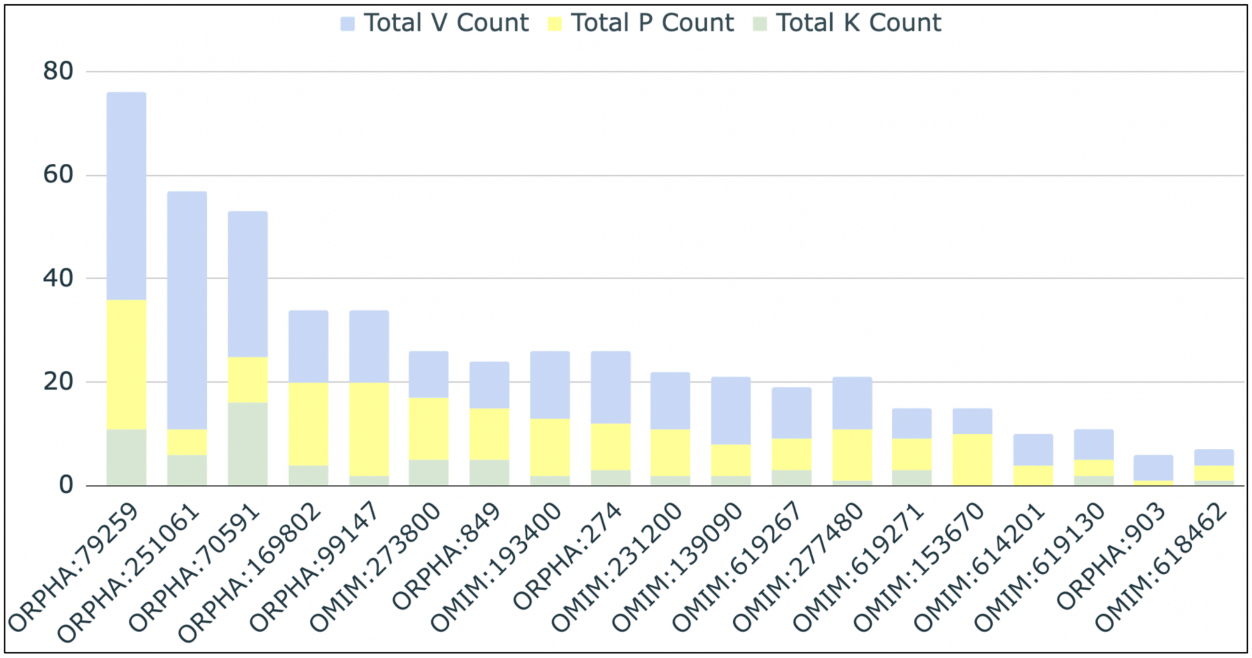


**Supplementary Pilot Study Fig.1**: The barplot depicts the total count of *V, P* and *K* phenotypes across all the 19 diseases associated with "Abnormality of von Willebrand factor".

Sanskrit shloka from original text with description of bleeding related phenotypes due to *Pitta* perturbation.

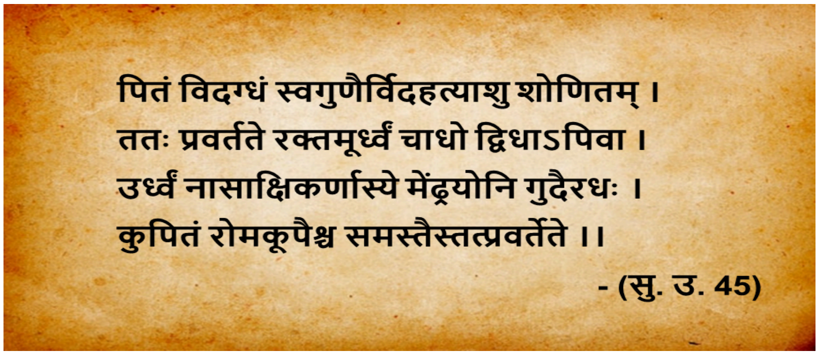


When the *Pitta* dosha gets vitiated by its own qualities, it affects the blood quickly in the body. This aggravated *pitta* mixed blood circulates in the body in an upward and downward direction. Upwards, it causes bleeding through the nose, eyes, ears, mouth; and downwards, it causes bleeding through the genitals (menstrual passage in women), anus in excessive aggravation, causes bleeding through all hair follicles of the body.

**Case Study 2- Associations of clinical features of *dosha* with modern diseases**

We carried out a similar exercise starting from a clinical condition of Ataxia (HP:0001251). Ataxia HPO yields 1255 diseases along with a set of 4029 non-redundant phenotypes (HPO version: May version, dated 2022-10-05). Despite its heterogeneity, an Ayurveda clinician in practice would ascribe ataxia as a disease of *Vata* manifestation (1). A similar exercise as above was conducted on a set of 25 diseases associated with ataxia. As is evident, in contrast to the bleeding associated diseases, this set had a more frequent and significant presence of *V* features (Supplementary Pilot Study Fig.2).

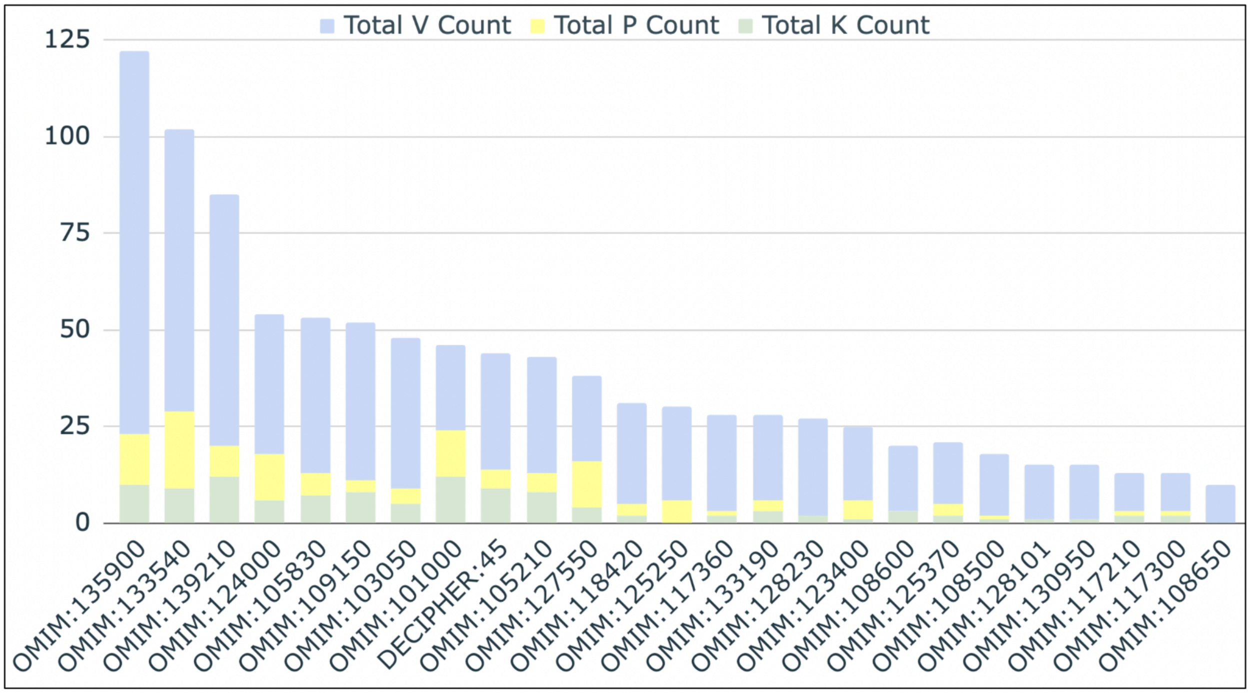


**Supplementary Pilot Study** **<u>Fig. 2:</u>** The barplot depicts the total count of *V*, *P* and *K* phenotypes across 25 diseases (subset) associated with "Ataxia". The cumulative count of V is observed high for this subset.

**Case Study 3 - Associations of clinical features of *dosha* with modern diseases**

Ayurveda also has descriptions of *dosha* specific cellular functions that have been correlated in our earlier studies (5). For example *Vata* is described in Ayurveda to be responsible for cell division and morphogenesis. Enhanced rates of cell proliferation rates have been reported in the transcriptome as well as cell lines derived from *Vata* individuals. (Cell Cycle). A query for "abnormality of cell cycle" in HPO provides four diseases associated with Fanconi anemia (Supplementary Pilot Study Table 5). Annotation of Fanconi anemia with Ayurveda *doshas* reveal enrichment of *Vata* (Supplementary Pilot Study Table 6, Supplementary Pilot Study Fig. 3)

**<u>Supplementary Pilot Study Table 5:</u>** List of syndromes with a search for "abnormality of cell cycle" term in the HPO database

| HPO Disease Id | HPO Disease Name |
|---|---|
| OMIM:227650 | Fanconi anemia |

| | |
|---|---|
| OMIM:227645 | Fanconi anemia, complementation group C |
| OMIM:227646 | Fanconi anemia, complementation group D2 |
| OMIM:600901 | Fanconi anemia, complementation group E |

**<u>Supplementary Pilot Study Table 6:</u>** Labeling of the phenotypes in one of the disease

| **HPO Disease Id** | **HPO Disease Name** | **HPO Term Id** | **HPO Term Name** | ***VPK* Mapping** |
|---|---|---|---|---|
| OMIM:227650 | Fanconi Anemia | HP:0030680 | Abnormal cardiovascular system morphology | *V* |
| | | HP:0001627 | Abnormal heart morphology | *V* |
| | | HP:0012210 | Abnormal renal morphology | *V* |
| | | HP:0001000 | Abnormality of skin pigmentation | *P* |
| | | HP:0003974 | Absent radius | *V* |
| | | HP:0009777 | Absent thumb | *V* |
| | | HP:0001903 | Anemia | *P* |
| | | HP:0001017 | Anemic pallor | *P* |
| | | HP:0000007 | Autosomal recessive inheritance | *V* |
| | | HP:0000978 | Bruising susceptibility | *P* |
| | | HP:0000957 | Cafe-au-lait spot | *V/P* |
| | | HP:0003221 | Chromosomal breakage induced by crosslinking agents | *V* |
| | | HP:0009943 | Complete duplication of thumb phalanx | *V* |
| | | HP:0000028 | Cryptorchidism | *V* |
| | | HP:0003213 | Deficient excision of UV-induced pyrimidine dimers in DNA | *V* |
| | | HP:0000081 | Duplicated collecting system | *K* |
| | | HP:0000086 | Ectopic kidney | *V* |
| | | HP:0000365 | Hearing impairment | *V/K* |
| | | HP:0000085 | Horseshoe kidney | *V* |
| | | HP:0000815 | Hypergonadotropic hypogonadism | *V* |
| | | HP:0001249 | Intellectual disability | *P* |
| | | HP:0001909 | Leukemia | *V/P/K* |
| | | HP:0003251 | Male infertility | *V* |
| | | HP:0000252 | Microcephaly | *V* |
| | | HP:0000568 | Microphthalmia | *V* |
| | | HP:0001875 | Neutropenia | *V* |
| | | HP:0001876 | Pancytopenia | *V* |
| | | HP:0003214 | Prolonged G2 phase of cell cycle | *V* |
| | | HP:0000104 | Renal agenesis | *V* |
| | | HP:0001896 | Reticulocytopenia | *V* |
| | | HP:0004322 | Short stature | *V/nV* |
| | | HP:0009778 | Short thumb | *V* |

| | | | | |
|---|---|---|---|---|
| | | HP:0001518 | Small for gestational age | *V* |
| | | HP:0000486 | Strabismus | *V* |
| | | HP:0001873 | Thrombocytopenia | *V* |

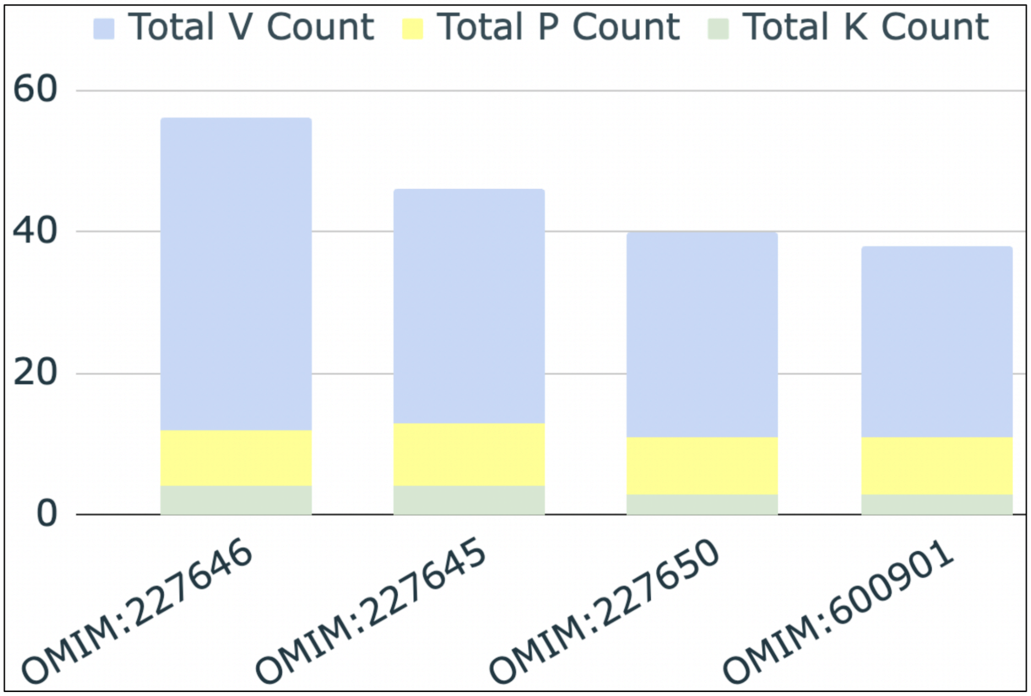


**Supplementary Pilot Study Fig.3:** The barplot depicts the total count of *V, P* and *K* phenotypes across all the 4 diseases. The cumulative count for *V* as a major phenotype is evident from the figure. Since cell division and morphogenesis is associated with imbalance of *V*

Based on the above exercise we surmised that there is a possibility for integrating Ayurveda and modern medicine clinical descriptions using the interface of HPO to probe the ontological links. This could be either through clinical features from ayurveda or modern medicine as well as functional attributes described for dosha. An extensive effort was undertaken to assign ~12,000 diseases associated with a non redundant set of over 10,000 phenotypes to understand the rare diseases from Ayurveda perspective.

### 2. Text

**Textual description in Ayurveda related to manifestation of Ciliopathies**

The normal function of transport and motility is governed by *V* (see box 1), which when perturbed manifests the features related to *K* (see box 2).

**1. Normal *Vata* function**

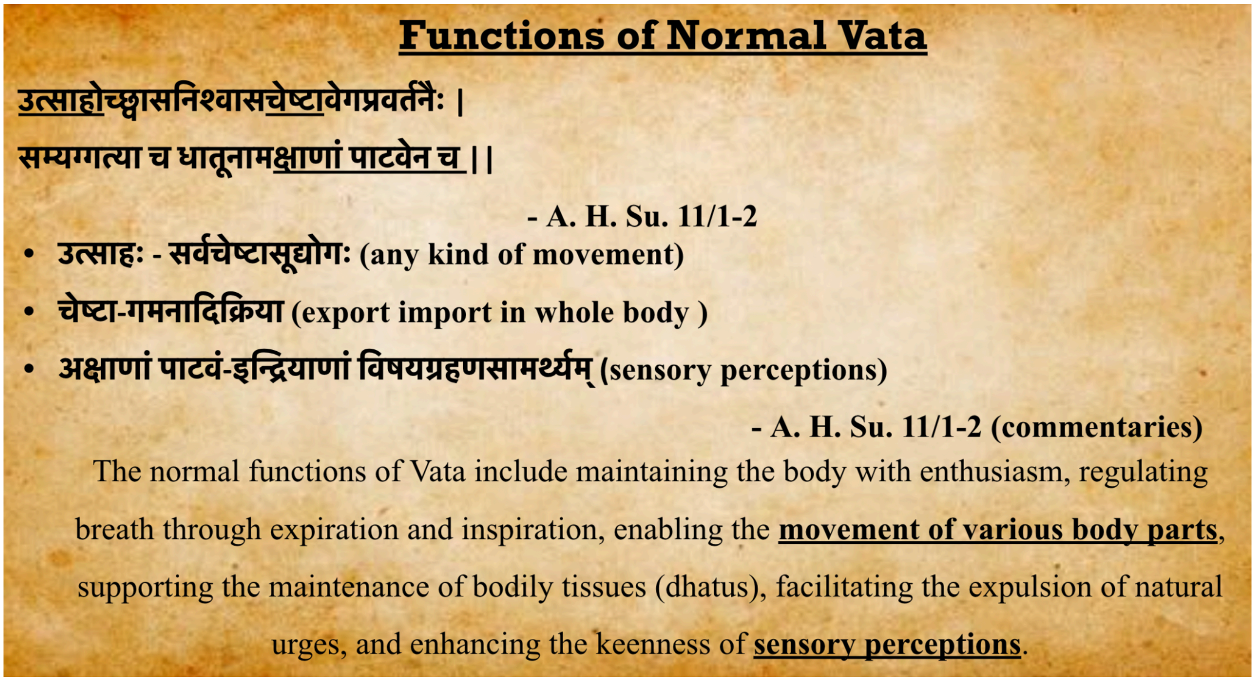


**2. When *Vata* function is perturbed**

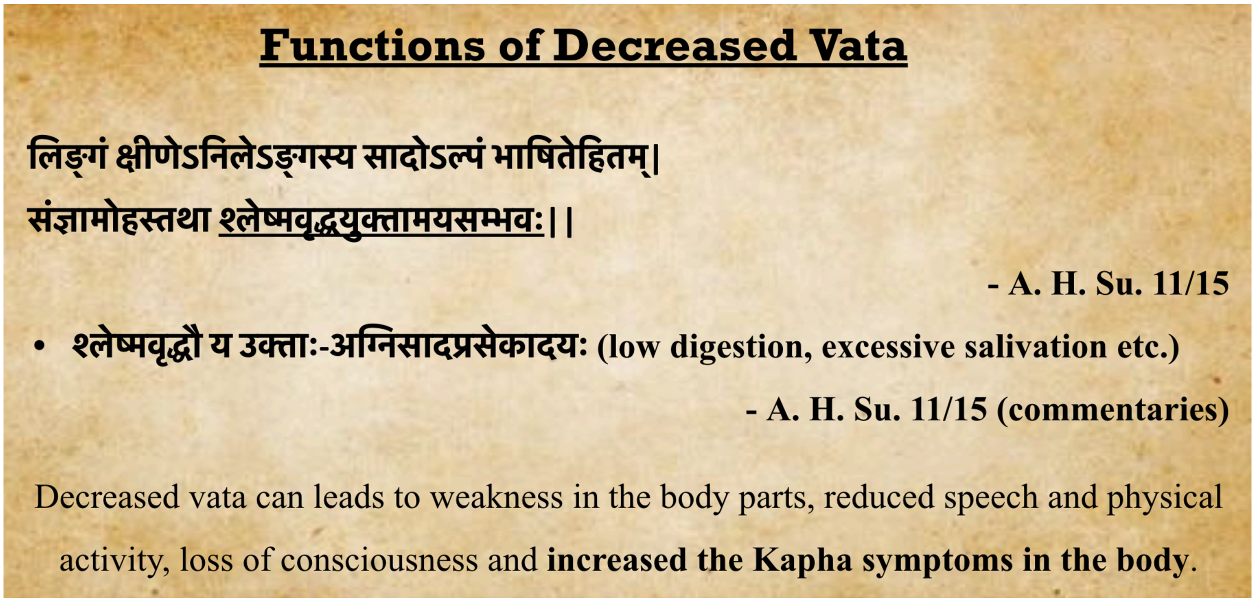


### 3. Methods

**Identification of clusters of diseases based on doshic proportions using Expectation Maximization (EM) algorithm**

Examination of the dataset revealed that the disease to HPO association is probabilistic in nature. The dataset shows that for a single disease, the associated features may or may not have been recorded for all the patients. This is quite understandable as not all patients who suffer from the same disease report all the symptoms, nor do doctors look for an exhaustive set of symptoms for a particular disease.

Expectation-Maximization (EM) clustering is a powerful algorithm used for probabilistic clustering, especially in situations where the data may comprise multiple underlying distributions. This algorithm iteratively finds local maximum likelihood parameters of the underlying distributions. The distributions may involve latent variables and unknown parameters which have to be inferred from given data observations. The method can support missing values in the data, which was highly suitable for our purpose. EM algorithm assigns a probability distribution to each instance which indicates the probability of it belonging to each of the clusters. The number of clusters may be specified a priori or determined by the algorithm through cross validation.

The key steps of the algorithm are explained below:

- **Initialization:**
    - The parameters of the Gaussian distributions i.e. the means, covariables and mixture weights are initialised with random values. The input value of the cluster can be given as an input or as in our case, the algorithm can find it iteratively through exploration and using the convergence properties.
- **Expectation Step (E-step):**

- For each data point xi, the probability ric , typically referred to as “responsibility” that it belongs to a Gaussian distribution c is computed using the multivariate Gaussian probability density function as follows:

$$r_{ic} = \frac{\pi_c N(x_i \mid \mu_c, \Sigma_c)}{\Sigma_{k=1}^{K} \pi_k N(x_i \mid \mu_k, \Sigma_k)}$$

  where K is the total number of clusters, πc is the mixing coefficient of the weight for the Gaussian distribution c, which was initialised in the earlier step, and N(x|μ, **Σ**) describes the probability density function (PDF) of a Gaussian distribution with mean μ and covariance **Σ**, with respect to datapoint x. N(x|μ, **Σ**) is computed as given below:

$$N(x_i, \mu_c, \Sigma_c) \; = \; \frac{1}{(2\pi)^{\frac{n}{2}} |\Sigma_c|^{\frac{1}{2}}} exp(-\frac{1}{2}(x_i - \mu_c)^T \Sigma_c^{-1} (x_i - \mu_c))$$

  The responsibility measures how much the c-th Gaussian distribution is responsible for generating the i-th data point.

- **Maximization Step (M-step):**

  In the M-step, the algorithm uses the responsibilities of the Gaussian distributions computed earlier to update the estimates of the model’s parameters.

- The weights πc, the means μc and the covariance **Σ**c are updated using the following equations -

$$\pi_c = \frac{\Sigma_{i=1}^{m} r_{ic}}{m}$$

$$\Sigma_c = \frac{\Sigma_{i=1}^{m} r_{ic} (x_i - \mu_c)^2}{\Sigma_{i=1}^{m} r_{ic}}$$

- The updated estimate is used in the next E-step to compute new responsibilities for the data points.

- The E-step and M-step are iteratively repeated till either a convergence or maximum number of iterations is reached.

- **Checking for Convergence**
  - Convergence is checked by evaluating the change in log-likelihood of the data, using the equation given below:

$$L(\Theta) = i\ k\ \ r_{ik} \log(\pi_c * N(x_i \mid \mu_k , \Sigma_k))$$

If the change falls below a specified threshold then it is said to converge. Once convergence is reached, the final parameters of the Gaussian mixture model are accepted as estimates of the underlying distributions. We have used the Weka package for implementing the EM algorithm. The exercise was repeated thrice with different initializations.

### 4. Figures

**Fig S1(a):** The modules represented belong to the cluster C1: This cluster is a mix type with sharing characteristics from *Vata* and *Pitta*. The processes shown here includes immune response, homeostasis and transport of small molecules

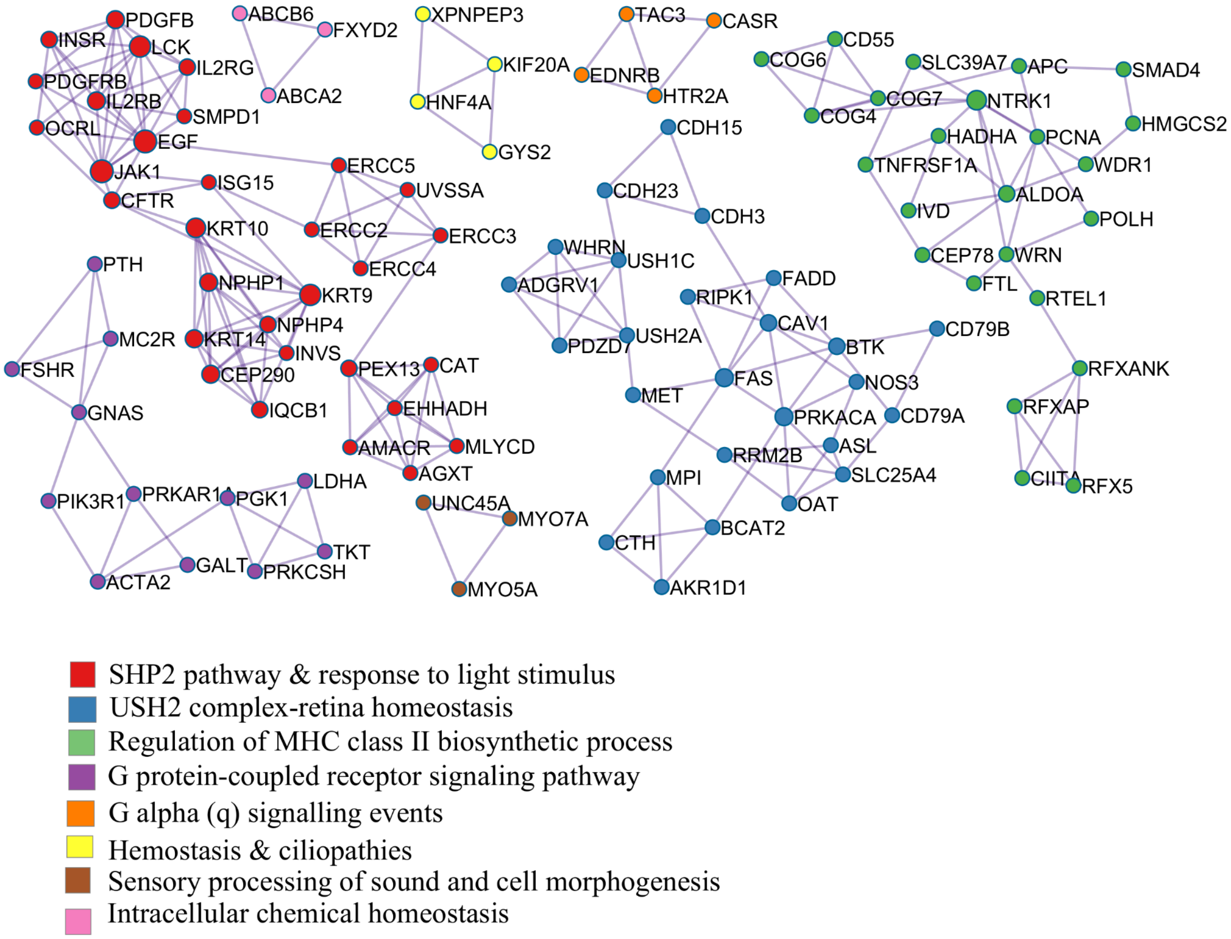

**Fig S1(b):** The modules represented belong to the cluster C2: This cluster is dominated with characteristic *Pitta* features. The processes observed here majorly include immune and inflammation along with cell activation and signaling.

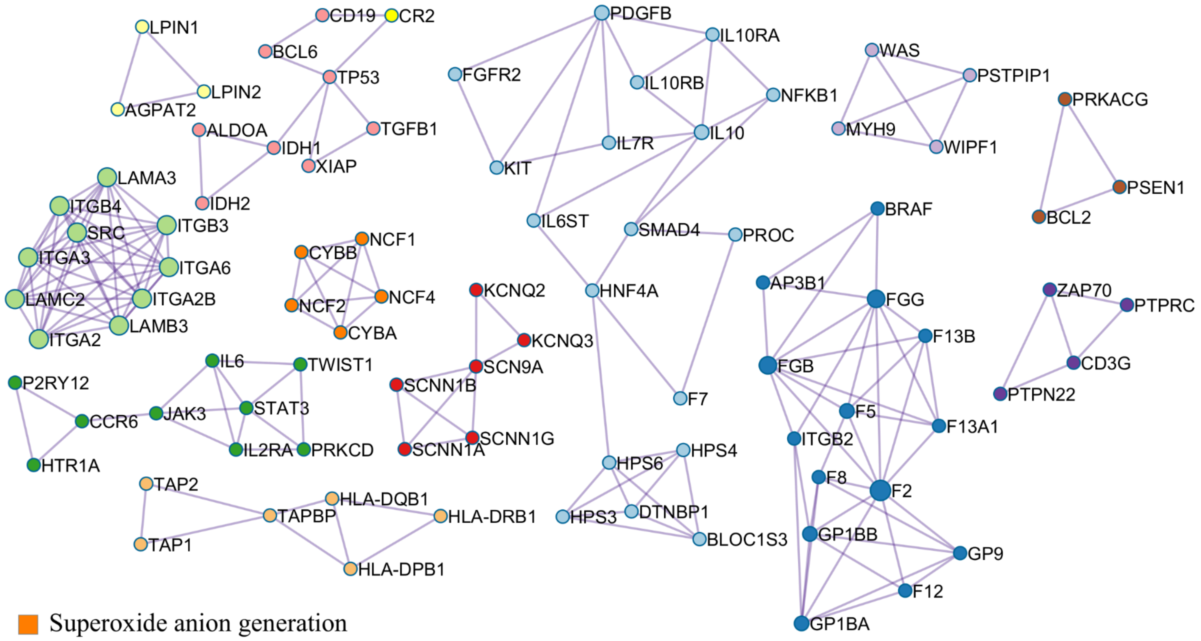


Superoxide anion generation
Parasitic Infection Pathways
T cell receptor signaling pathway
Triglyceride biosynthetic process
Hemopoiesis
Platelet dense granule organization
Blood coagulation, fibrin clot formation
Focal adhesion
Cell activation- interleukin-2-mediated signaling pathway
Lymphocyte differentiation
Sensory perception of salty taste
Antigen processing and presentation of peptide antigen

**Fig S1(c):** The modules represent the cluster C3: This cluster is dominantly *Pitta* with a small share of features from *Vata*. The processes here include inflammation and telomere activity.

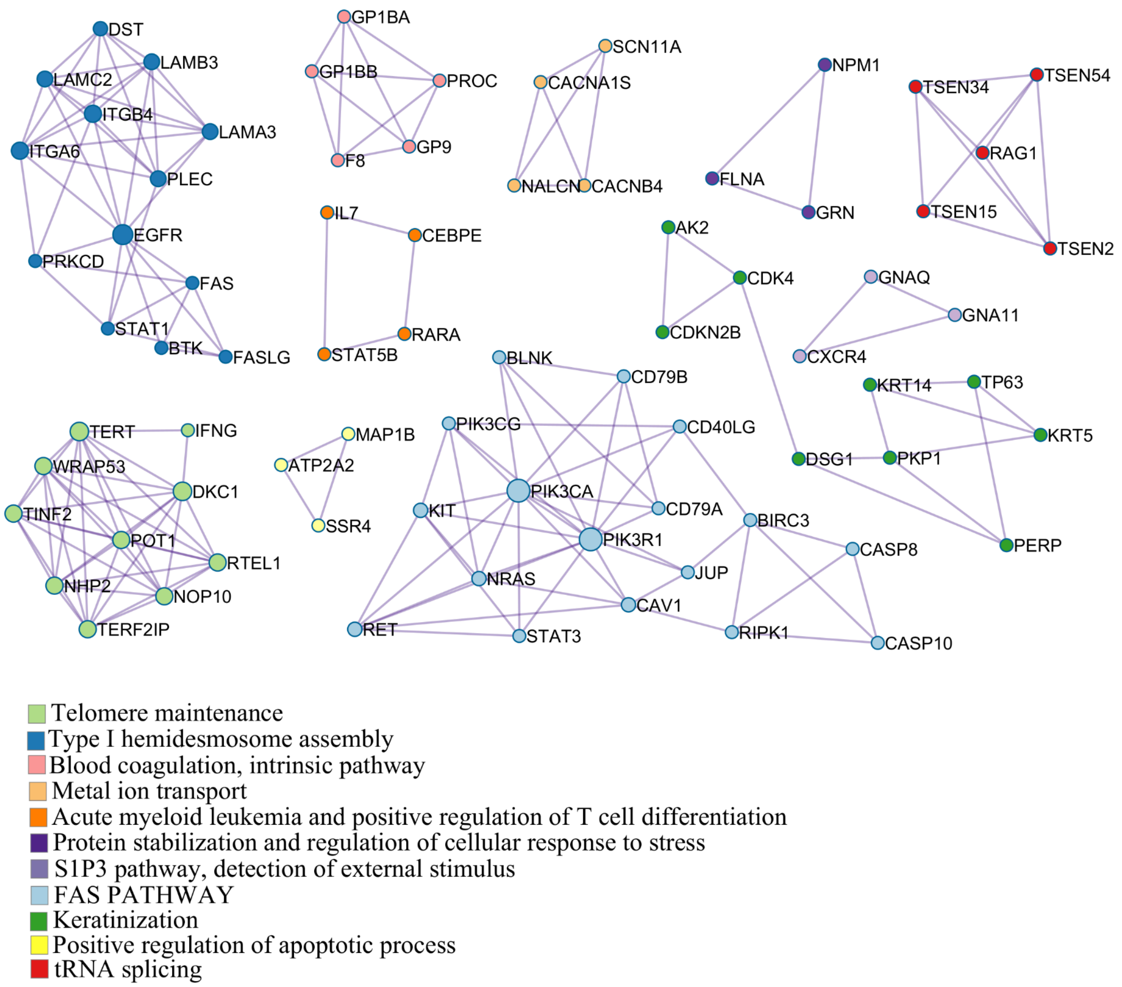

**Fig S1(d):** The modules represent the cluster C4. The cluster is dominantly *kapha*. Hence the modules are  associated with ciliopathies, visual perception and metabolism.

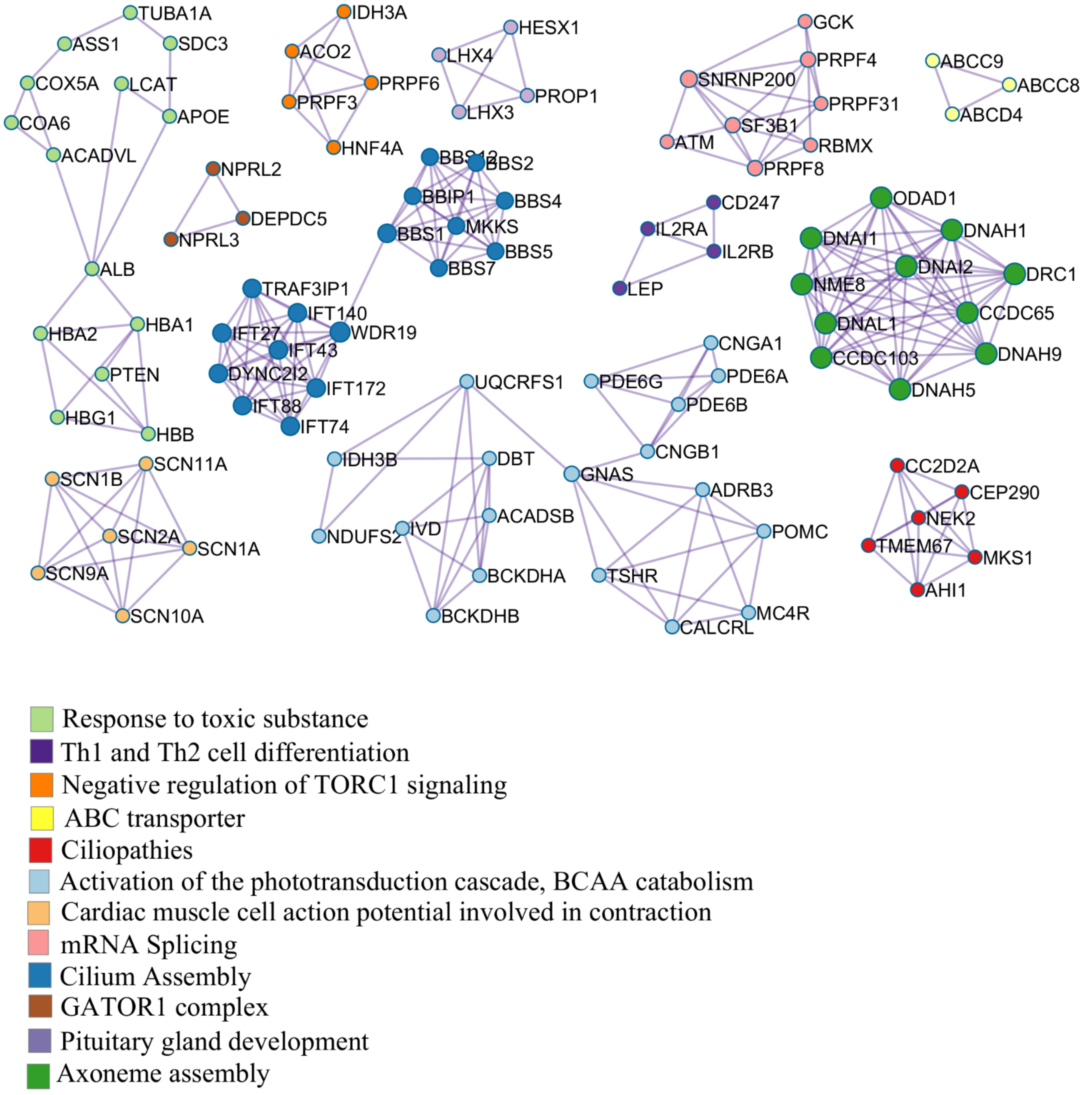

**Fig S1(e):** The modules represented are from cluster C5. This cluster is highly rich with characteristics of *Vata*. The modules for this cluster are distinct from other clusters sharing characteristics of *Vata*. The distinct processes associated with this cluster are DNA damage, DNA damage response, chromatin organization and regulation of cell cycle.

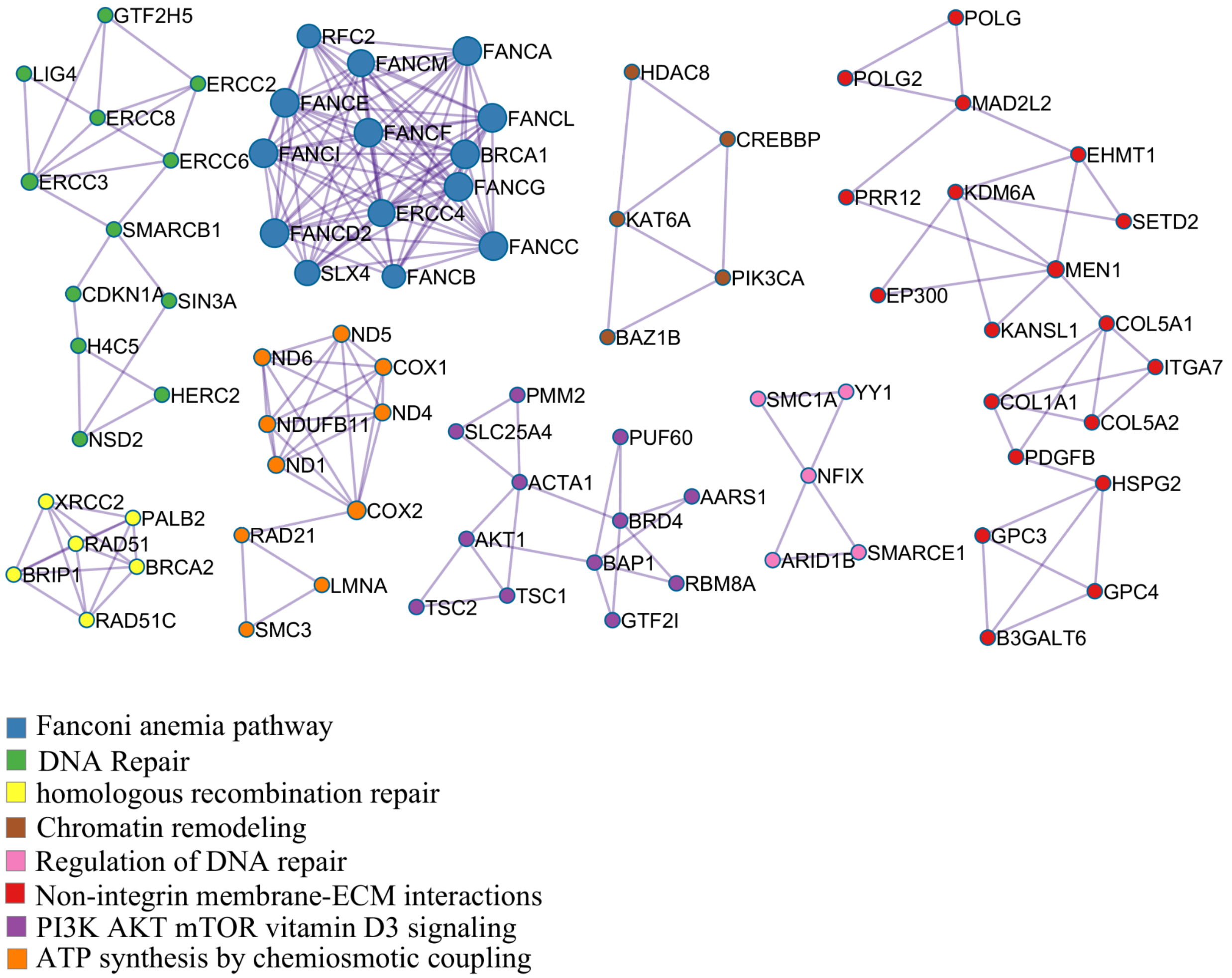

**Fig S2:** This figure shows the network for all ciliary genes from all the clusters, this contains 5 major hubs of pathways most of them are dominated by Cluster C4 & C0**.**

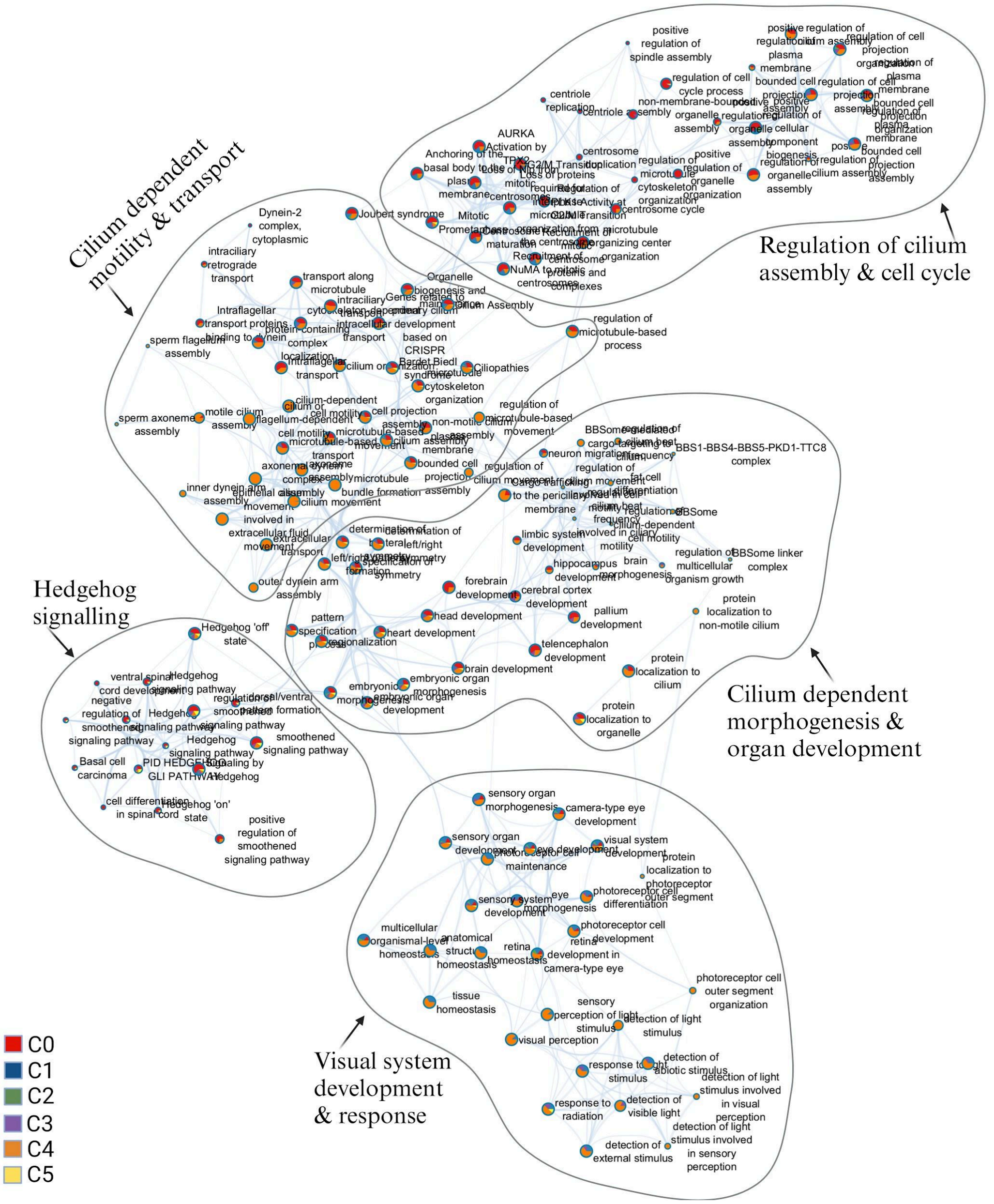

**Fig S3(a):** This figure shows the contribution of ciliary and remaining genes from cluster C4 in cellular pathways. The network has two inter-connected hubs, one cilium assembly & functions, which is dominated by overlapping ciliary genes, and other cellular responses to biomolecules dominated by non- overlapping genes.

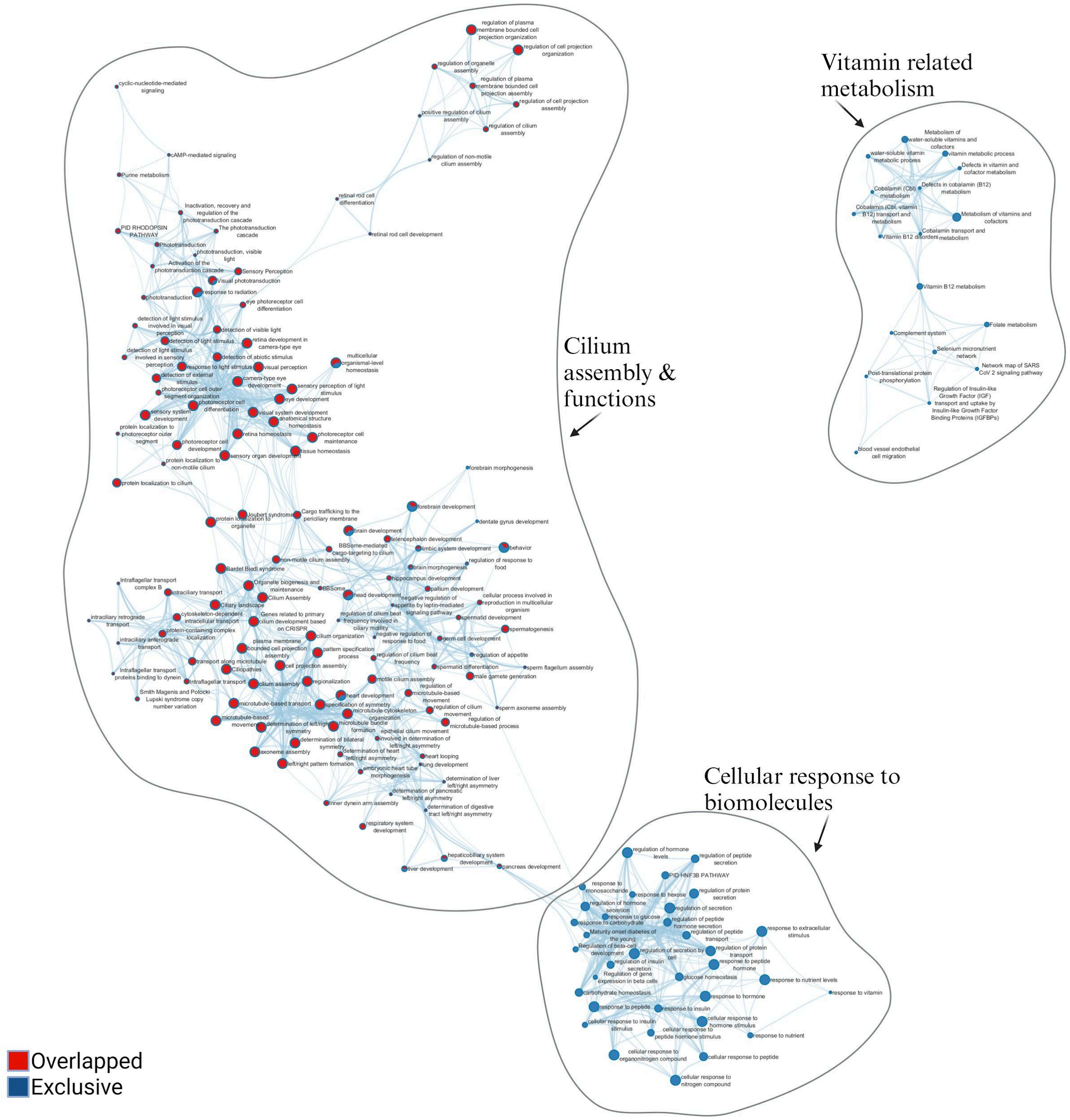

**Fig S3(b):** A network analysis of cluster C0 overlapping ciliary known genes & non-overlapped both, showing the interconnectedness between cilium-related signalling and morphogenesis & organ development, dominated by ciliary genes and non-overlapped ones, respectively.

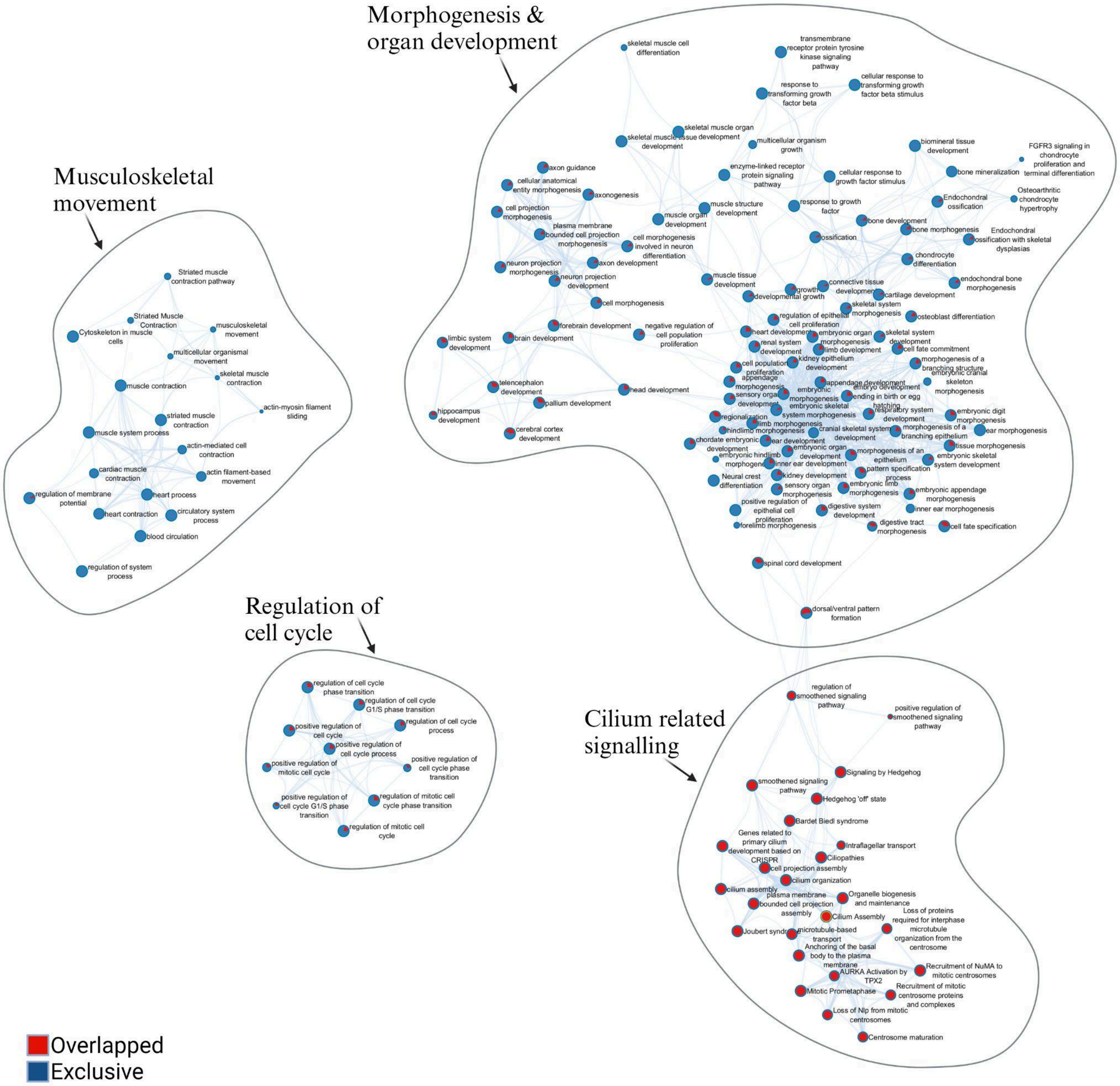

**Fig S4:** Network analysis of cluster C5. Three clusters i.e. C0, C1and C3 have been observed with characteristic *Vata* features in share with *Pitta* features. Cluster C5 is dominant in *Vata* features involved in distinct processes like DNA damage, DNA damage response, chromatin organization and regulation of cell cycle, which have been found absent in other three clusters.

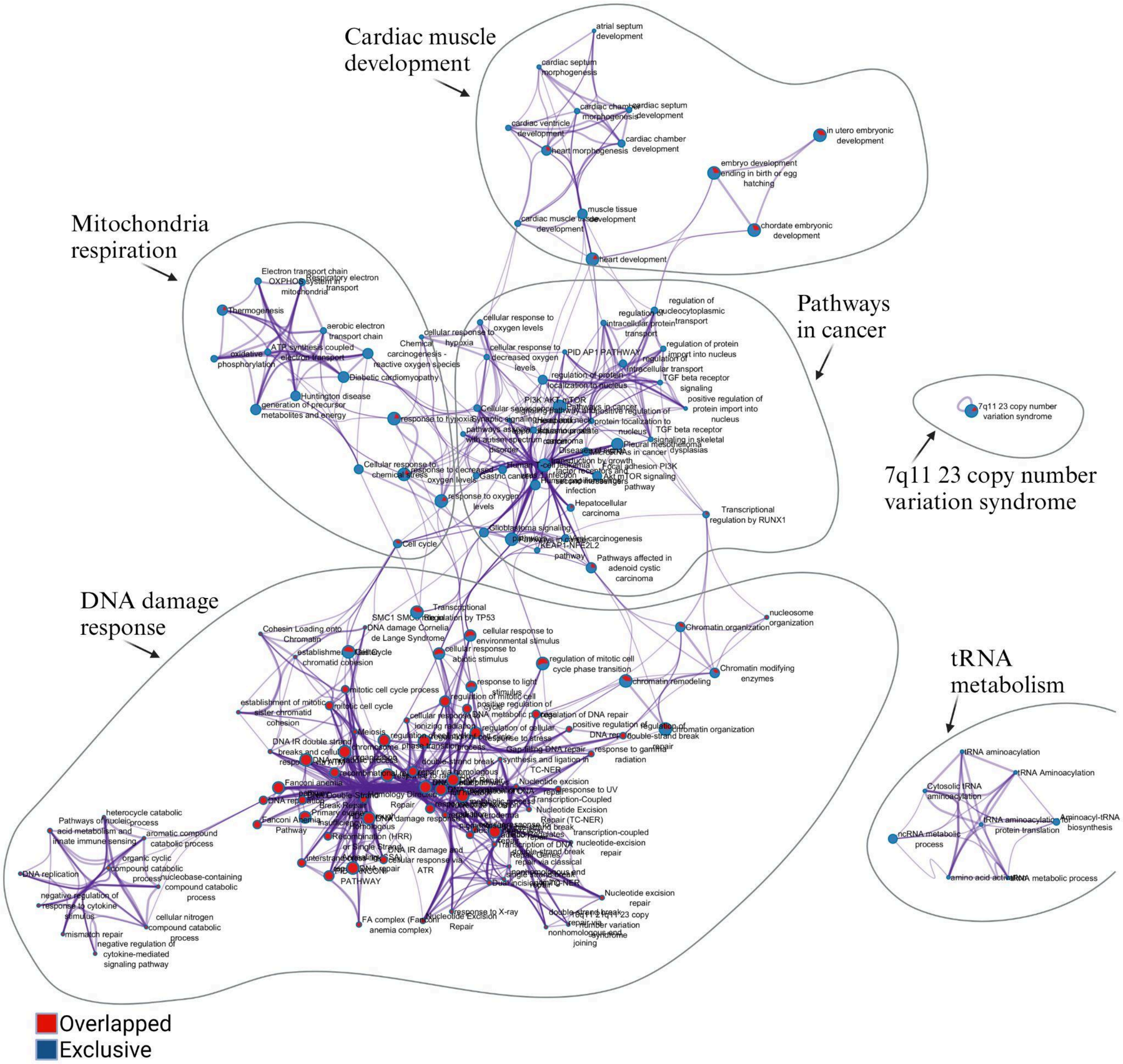
